# Single-Cell and Spatial Transcriptomic Analyses Deciphering the Three-Layer Architecture of Human Tuberculosis Granulomas

**DOI:** 10.1101/2024.07.15.603490

**Authors:** Xia Yu, Jie Wang, Peihan Wang, Xiaoqiang Liu, Cuidan Li, Yingjiao Ju, Sitong Liu, Yujie Dong, Jing Wang, Bahetibieke Tuohetaerbaike, Hao Wen, Wenbao Zhang, Haitao Niu, Sihong Xu, Chunlai Jiang, Xiaoyi Jiang, Jing Wu, Hairong Huang, Fei Chen

**Author notes:** These authors contributed equally to this work* (Xia Yu, Jie Wang, Peihan Wang, Xiaoqiang Liu, Cuidan Li, Yingjiao Ju). Correspondence:* (Fei Chen); (Hairong Huang).

## Abstract

**Background:** Granulomas (defining tuberculosis histopathological feature) are central to the host’s defense against *Mycobacterium tuberculosis*, critically influencing patient outcomes. However, knowledge of human granulomas’ structure and function are incomplete. This study employs single-cell and spatial transcriptomics to dissect human granuloma’s cellular composition, structure, communication and function from 19 pulmonary, lymphatic and skeletal samples.

**Results:** Our study identified nine key immune-activated/signaling-active cell clusters. Notably, we delineated a three-layered granuloma structure: a core with macrophages (Macro-c09, Macro-c10) and occasional fibroblasts (Fib-c03); a fibroblast-rich (Fib-c01) periphery; and an immune-infiltrated intermediate layer comprising diverse immune-cells recruited by strong signaling-molecules (SPP1/MIF) from core/periphery cells. This study also shows granuloma heterogeneity across individuals and tissues.

**Conclusions:** By merging scRNA-seq with ST-seq, we offer an intricate single-cell perspective of granulomas’ spatial-structure and formation mechanisms, identify signaling-molecules and significantly changed genes as potential targets for host-directed tuberculosis immunotherapy, highlight fibroblasts’ crucial role in granuloma formation, and provide an important reference/improved understanding of TB.

## Background

Tuberculosis (TB), caused by *Mycobacterium tuberculosis* (*Mtb*), was historically the most lethal infectious disease until COVID-19 (*1*). Granulomas, the most defining morphological and histopathological characteristic lesions in TB, are the primary arenas of host defense against *Mtb* in the lungs and other organs (*2*). These structures maintain a delicate balance between pro-inflammatory and anti-inflammatory responses, crucial for patient outcomes. Granulomas can function as protective barriers, entrapping and even eliminating *Mtb* with the collective effort of immune cells like macrophages (*3*). Yet, *Mtb* may exploit the granuloma’s closed, anti-inflammatory and lipid-rich environment to survive, proliferate, and eventually disseminate (*4*), leading to severe disease progression. Thus, the TB clinical outcomes largely depend on this delicate balance, underscoring the importance of granuloma analysis.

Histological studies describe granulomas as structured immune cell aggregates with a macrophage-rich core and a periphery of lymphocytes and fibroblasts (*5, 6*). Recent single-cell and spatial technologies have advanced our understanding but have limitations. To date, three animal-based single-cell RNA sequencing (scRNA-seq) studies (zebrafish (*7*), rhesus macaque (*8*), and cynomolgus macaque (*9*)) have been insightful but cannot wholly mirror human responses. Three studies focused on granulomas’ spatial structure have been performed so far. One mouse lung granuloma study utilized in situ sequencing with 34 immune markers, delineating increased transcriptomic co-localization networks during infection progression and significantly enrichment of certain activated macrophage subtypes (*10*). The other two studies have harnessed spatial omics to analyze human TB granulomas with limited gene markers. One employed MIBI-TOF with 37 gene markers, and outlined 19 cell subsets across several tissues (*11*). Another employed spatial mapping approach using four gene markers, revealing enriched leukocyte around necrotizing granulomas (*12*). Overall, most single-cell and spatial studies utilize animal granuloma models, which do not fully replicate human disease dynamics, and the only two human studies have been restricted by limited genetic markers, offering only a glimpse of the complicate immune-environment.

Our study employs single-cell and spatial transcriptomics to dissect human TB granulomas from 19 pulmonary, lymphatic and skeletal lesion samples. By integrating scRNA-seq with spatial-seq, we identify nine key cell clusters and delineate a comprehensive cellular landscape of human granulomas, highlighting their consistency and heterogeneity. We describe a consistent three-layer spatial cellular architecture, with similar cell composition, functions and intercellular communications, but also uncover heterogeneity linked to disease severity. Overall, our study’s uniquely highlights the intricate three-layer structure of human granulomas, offering new insights into their formation and response to *Mtb*.

## Results

### Single-cell transcriptomic landscape of TB granulomas in pulmonary, lymphatic and skeletal lesion samples

To characterize the single-cell landscape of TB granulomas across various tissues, we performed scRNA-seq analysis of 11 samples from three pulmonary, five lymphatic, and three skeletal TB patients using the 10x Genomics platform (table S1; Fig. 1A). After filtering, 111,967 cells were retained across the three tissue types: pulmonary (32,200 cells), lymphatic (49,409 cells), and skeletal (30,358 cells). These cells were categorized into myeloid, lymphoid, and stromal cell populations, and further into 15 major cell types (fig. S1A, B). In all 11 samples, macrophages (9,743, 8.70%) dominated among myeloid cells, CD4+ T cells (37,173, 33.20%), CD8+ T cells (22,261, 19.88%), and B cells (18,092, 16.16%) were prevalent within lymphoid cells, and fibroblasts (3,580, 3.20%) were the predominant stromal cell type (fig. S1C). Our detailed analysis focused on these five cell types: macrophages, CD4+ T cells, CD8+ T cells, B cells, and fibroblasts. Further re-clustering revealed 71 distinct cell clusters, underscoring the diversity and heterogeneity within and across tissues (Fig. 1B-H).

**Fig. 1.**
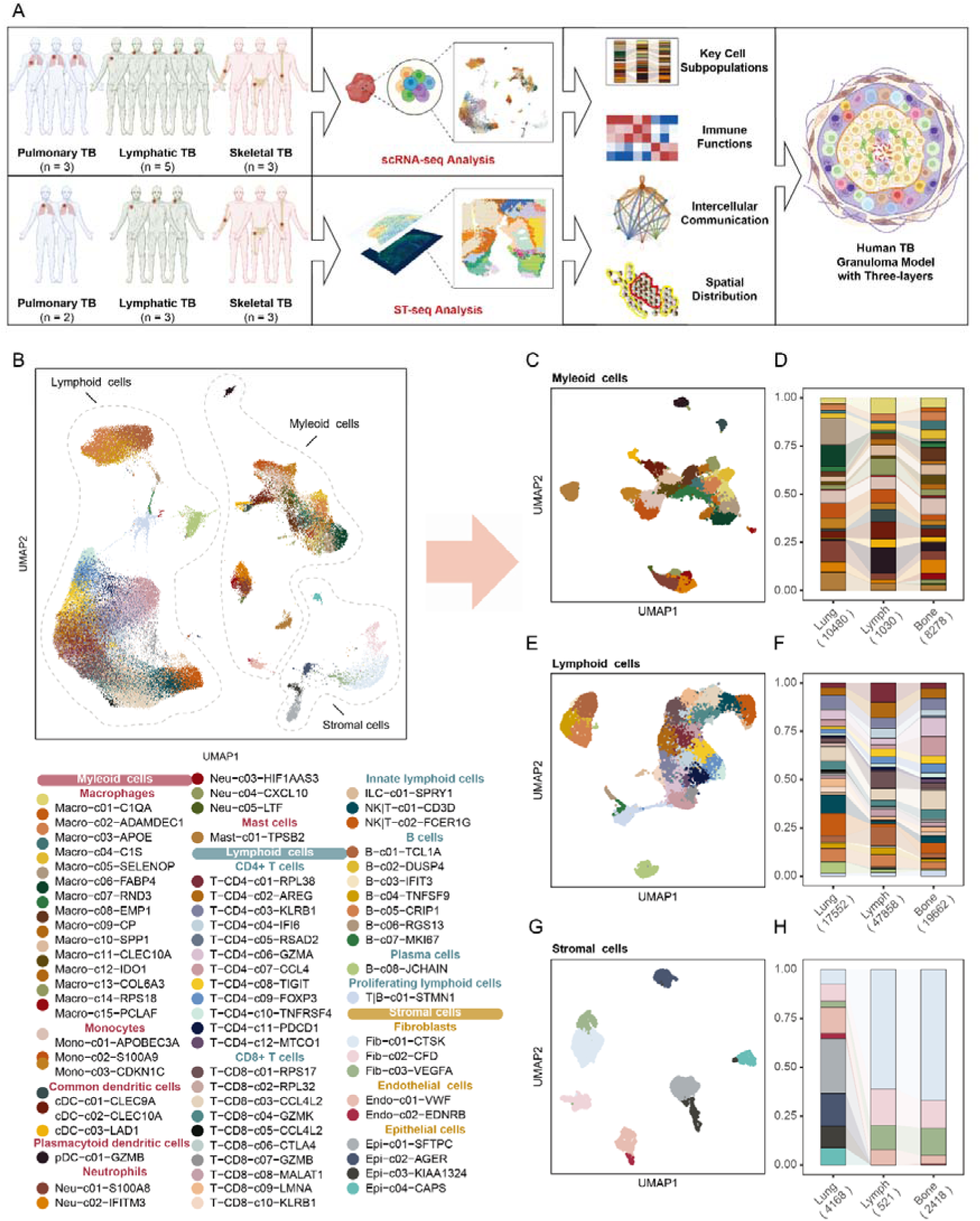
Single-cell and spatial transcriptomics analyses of TB granulomas in pulmonary, lymphatic, and skeletal Tissues. **(A)** Study overview: Tissue specimens collected from patients with pulmonary, lymphatic, and skeletal tuberculosis were subjected to scRNA-seq and ST-seq analysis, elucidating the key cell subpopulations, immune function, and intercellular communication and spatial distribution, thereby inferring the three-layer spatial architecture of human TB granulomas (Created with BioRender.com). **(B)** UMAP Plot of 111,967 cells colored according to their respective identities among 71 distinct cell subpopulations. **(C, E, and G)** UMAP Plots of (C) myeloid, (D) lymphoid, and (E) stromal cell subpopulations. **(D, F, and H)** Bar Plots of the proportions of (C) myeloid, (D) lymphoid, and (E) stromal cell subpopulations in pulmonary, lymphatic, and skeletal tissues.

**Macrophages**, serving as the primary host cells for *Mtb*, are predominant among myeloid cells and divided into 15 distinct cell clusters (Fig. 2A and fig. S2A, B). For each cluster, we conducted Gene Set Enrichment Analysis (GSEA) with gene sets from the Molecular Signatures Database (MsigDB) Hallmark collection, GO knowledgebase, and KEGG database to determine their immune-related characteristics and potential pro-inflammatory roles. Additionally, we performed cell-cell communication analysis to assess the potential strength of intercellular communication among the clusters. Notably, four cell clusters – Macro-c09, Macro-c10, Macro-c12 and Macro-c08 – stand out for their strong pro-inflammatory responses and/or intercellular communication (Fig. 2B, C).

**Fig. 2.**
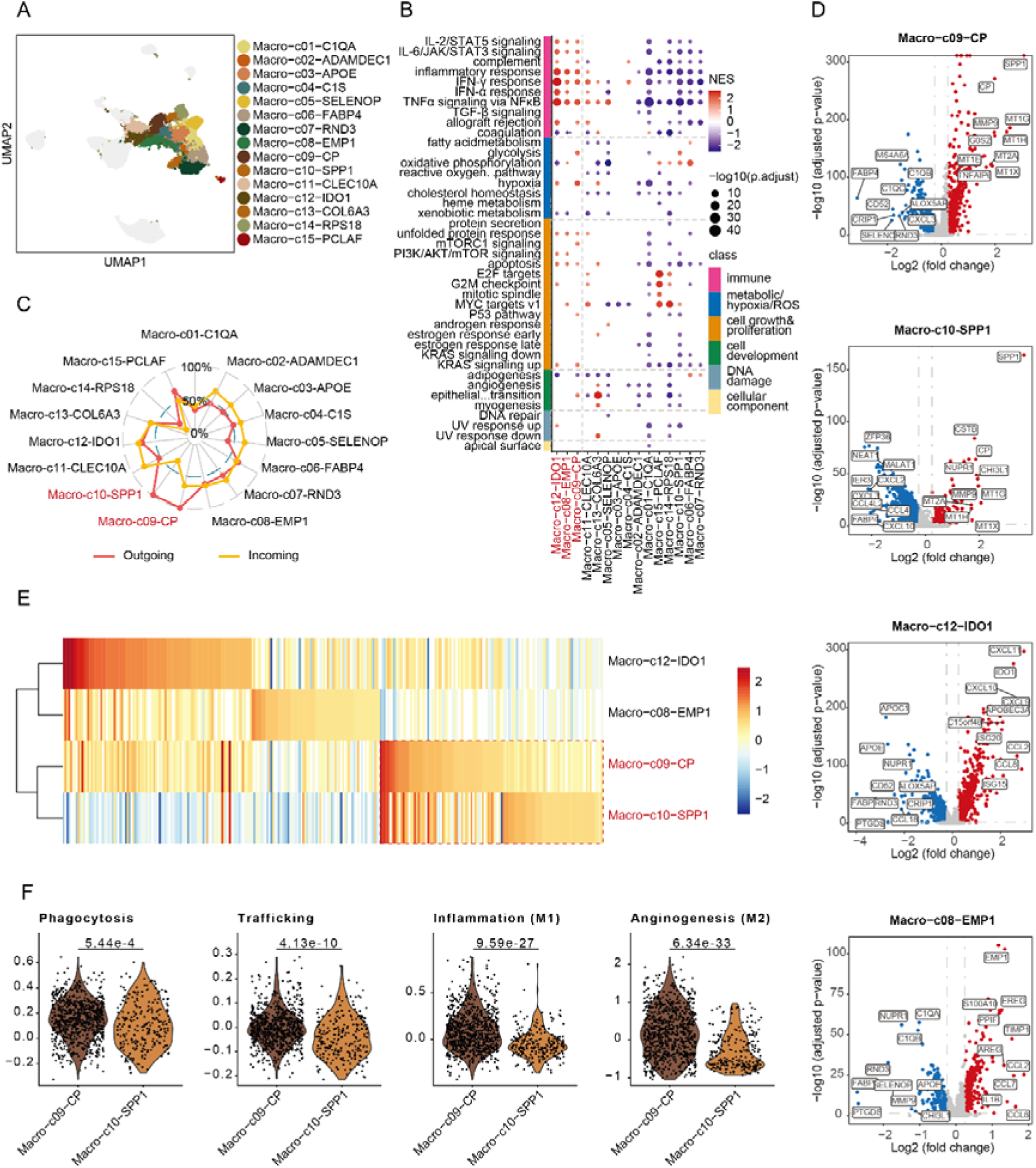
Macrophage subpopulations of TB granuloma samples. **(A)** UMAP plot of macrophage cells across all samples colored according to the identities of cell subpopulations. **(B)** GSEA analyses for macrophage subpopulations using MSigDB hallmark gene sets. Color intensity represents normalized enrichment score (NES), while dot size reflects statistical significance. **(C)** Radar plot showing intercellular interaction of macrophage subpopulations categorized as outgoing and incoming signaling. **(D)** Volcano plots showing DEGs in Macro-c12, Macro-C08, Macro-c09 and Macro-c10 compared to other macrophage cells. Red and blue dots denote up-regulated and down-regulated DEGs, respectively. **(E)** Heatmap displaying the top10 DEGs in Macro-c12, Macro-C08, Macro-c09 and Macro-c10. **(F)** Violin plots showing the phagocytosis, trafficking, inflammation and anginogesis scores for Macro-c12, Macro-C08, Macro-c09 and Macro-c10.

Macro-c09 is unique for its combined strong pro-inflammatory activity and outgoing intercellular signaling (Fig. 2B, C). It displays significantly upregulated pro-inflammatory interleukin (IL)-1, toll-like receptor (TLR), and nod-like receptor (NLR) pathways (fig. S2C, D). Importantly, elevated levels of secreted phosphoprotein 1 (SPP1) and matrix metalloproteinase 9 (MMP9) link this cell cluster to collagen remodeling and fibrosis (*13*) (Fig. 2D), relevant to scar formation (*14*), and pulmonary fibrosis (*15*). Macro-c09 also exhibits a heightened cellular response to metal ions (fig. S2C), crucial for antimicrobial activity (*16, 17*), indicated by the expression of Metallothionein (MT) family genes (MT1G, MT1H, MT1X, MT2A, and MT1E) (*18, 19*) (Fig. 2D).

Macro-c10 is characterized by its potential for strong intercellular communication but lacks pro-inflammatory activity (Fig. 2B, C). Interestingly, Macro-c10 shows similar gene expression profile with Macro-c09, with adjacent positions on the UMAP plot (Fig. 2A) and a substantial overlap (seven) in their top 10 highly expressed genes (including SPP1, MMP9, and MT family genes) (Fig. 2D, E). Like Macro-c09, Macro-c10 too exhibited an enhanced cellular response to metal ions in GO analysis (fig. S2C). On the other hand, Macro-c10 showed weak pro-inflammatory function (Fig. 2B) and incoming signaling (Fig. 2C), as evidenced by diminished functions in key processes like phagocytosis, trafficking, inflammation (M1 feature), and angiogenesis (M2 feature; Fig. 2F) (*20*), suggesting a relatively inactive or potentially dysfunctional state for Macro-c10 compared to Macro-c09.

Macro-c12 is the most pro-inflammatory macrophage cluster with moderate intercellular communications (Fig. 2B, C). Another notable finding is the increased IDO1 expression in Macro-c12 (Fig. 2D), which has been previously observed in core macrophages of human TB granulomas (*11*).

Macro-c08 also exhibits strong pro-inflammatory properties with moderate intercellular communications (Fig. 2B, C). A significant increase in epithelial membrane protein 1 (EMP1) expression is noted in volcano plot (Fig. 2D), aligning with previous findings on epithelioid macrophages in TB granulomas (*7, 21*).

#### CD4+ T cells

Among lymphoid cells, CD4+ T cells are the most prominent, subdivided into 12 cell clusters (Fig. 3A, fig. S3A-C and table S2). Notably, the T-CD4-c07 effector T cell cluster exhibits the most significant enrichment of immune-related genes (Fig. 3B) and robust intercellular communication (both incoming and outgoing; Fig. 3C), highlighting its important role in immune defense against *Mtb* infection. RNA velocity-based cell trajectory analysis positions T-CD4-c07 at the end of T cell development, demonstrating that this potential pro-inflammatory CD4^+^ T cell cluster likely evolves from other T cells during *Mtb* infection (fig. S3B). This cell cluster also exhibits upregulation of pro-inflammatory genes, like IFNγ and some chemokines (Fig. 3D), suggesting its potential role in macrophage recruitment and activation to combat *Mtb* invasion (*22*). Interestingly, increased expression of two granzyme genes (GZMB and GZMA) suggests potential cytotoxicity (Fig. 3D), aligning with prior studies (*23, 24*). Further T cell receptor (TCR) analysis reveals notable clonal expansion within the T-CD4-c07, possibly crucial for recognizing TB antigens (Fig. 3E).

**Fig. 3.**
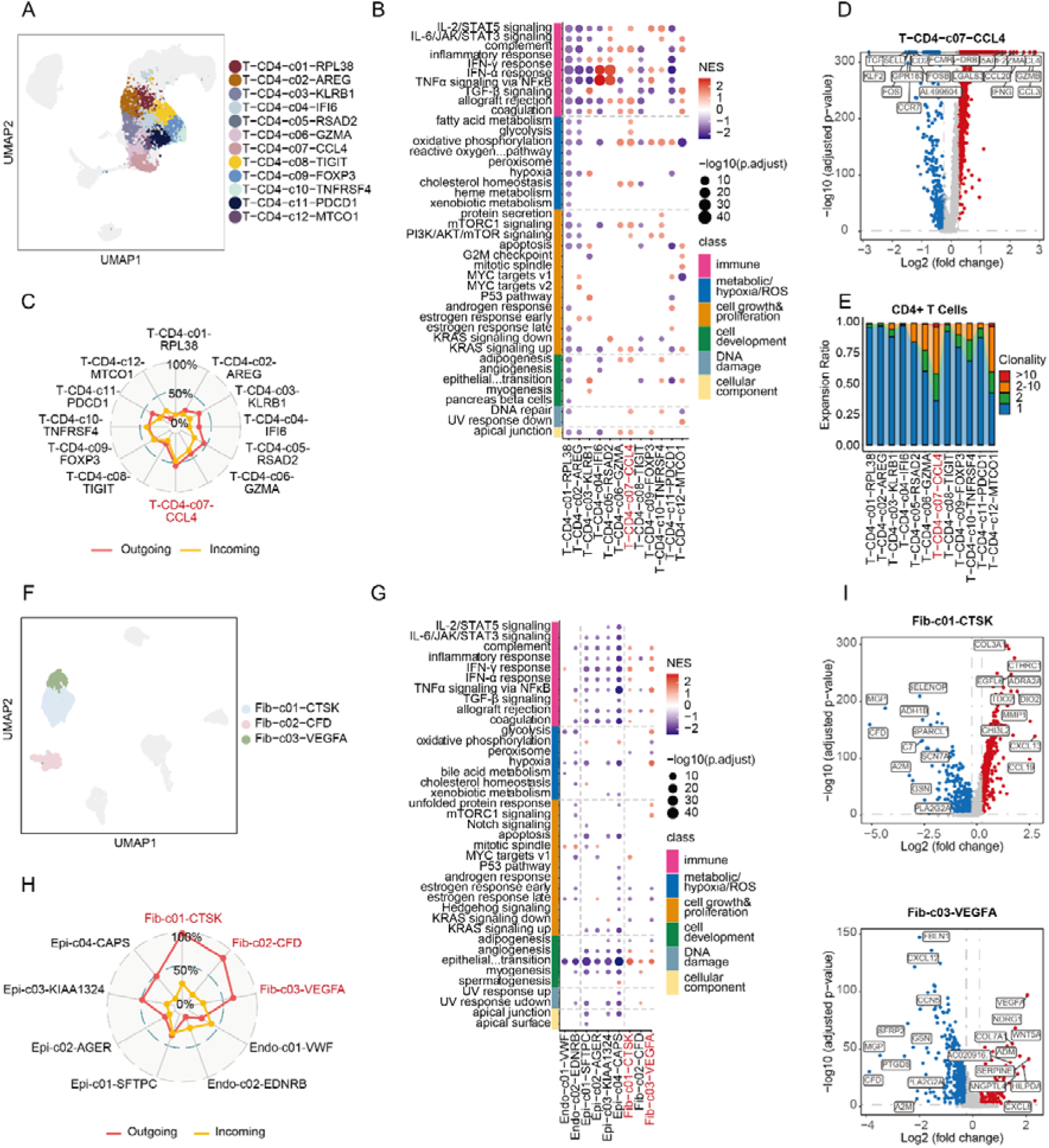
CD4^+^ T cell and fibroblast subpopulations of TB granuloma samples. **(A)** UMAP plot of CD4^+^ T cells across all samples colored according to the identities of cell subpopulations. **(B)** GSEA analyses for CD4^+^ T cell subpopulations using MSigDB hallmark gene sets. Color intensity represents NES, while dot size reflects statistical significance. **(C)** Radar Plot showing intercellular interaction of CD4^+^ T cell subpopulations categorized as outgoing and incoming signaling. **(D)** Volcano plots showing DEGs in T-CD4-c07 compared to other CD4^+^ T cells. Red and blue dots denote up-regulated and down-regulated DEGs, respectively. **(E)** TCR clonality in CD4^+^ T cell subpopulations. **(F)** UMAP plot of fibroblasts across all samples colored according to the identities of cell subpopulations. **(G)** GSEA analyses for stromal subpopulations using MSigDB hallmark gene sets. **(H)** Radar Plot showing intercellular interaction of stromal subpopulations categorized as outgoing and incoming signaling. **(I)** Volcano plots showing DEGs in Fib-c01 and Fib-c03 compared to other fibroblast cells. Red and blue dots denote up-regulated and down-regulated DEGs, respectively.

**CD8+ T cells** are second in prominence among lymphoid cells and re-clustered into 10 cell cluster (fig. S3D-F). Notably, the T-CD8-C07 effector T cell cluster displays strong pro-inflammatory features (like IFNγ and chemokines) (fig. S3G), intercellular communications (fig. S3H), cytotoxicity (including granzymes; fig. S3I), and clonal expansion (fig. S3J), emphasizing its important role in the immune defense against *Mtb*.

**B cells** are the third prevalent among lymphoid cells and re-clustered into nine cell clusters (fig. S4A-C). B-c08, the sole plasma cell cluster, displays heightened expression of various antibody genes but shows restricted pro-inflammatory activity and intercellular communication (fig. S4D, E), indicating its specialized role in humoral immunity over cellular immunity. This subpopulation also exhibits significant B cell receptor clonal expansion (fig. S4F, G), suggesting a potential for antibody production in reaction to *Mtb* infection (*25, 26*).

#### Fibroblasts

We identified 3,580 fibroblast cells and re-clustered them into three cell clusters (Fig. 3F and fig. S5A, B). When compared to other stromal cells, two fibroblast clusters, Fib-c01 and Fib-c03, showed relatively strong pro-inflammatory characteristics, including IFN responses and TNF signaling (Fig. 3G). Importantly, all identified fibroblast clusters showed strong outgoing intercellular signals, suggesting their roles in recruiting and activating immune cells (Fig. 3H). GO and KEGG also analyses showed a significant enrichment of chemokine- and cytokine-related pathways in Fib-c01 and Fib-c03, indicating their pro-inflammatory roles in activating and recruiting immune cells to sites of granuloma formation (Fig. 3I and fig. S5C, D). These findings align with similar observations in cancer research (*27*). Additionally, they showed enhanced responses to hyperoxia and oxidative stress (fig. S5C), processes known to tissue damage/fibrosis (*28*), further highlighting their involvement in TB granuloma formation.

### Spatial-seq reveals organized granulomas’ structure in pulmonary, lymphatic, and skeletal TB lesions

To investigate granuloma immune microenvironment, we performed ST-seq analysis on eight granuloma samples obtained from two pulmonary, three lymphatic and three skeletal TB patients (table S1). Two histopathologists verified each sample’s granulomas based on Hematoxylin and eosin (H&E) staining (fig. S6). Notably, granuloma-identified regions consistently presented more UMIs than adjacent non-granulomatous regions, indicating higher cell density in granulomas.

### Pulmonary granulomas show similar cell composition with some heterogeneity

We first performed UMAP analysis for unsupervised clustering of ST-seq data, classifying the two pulmonary lesion samples (T166390 and T167151) into 20 and 18 clusters, respectively. Pathological assessments identified two distinct granuloma clusters (g1 and g2) with differential gene expression profiles in each sample, reflecting the pulmonary granulomas’ heterogeneity, as previous research (*12, 29*) (Fig. 4A-C).

**Fig. 4.**
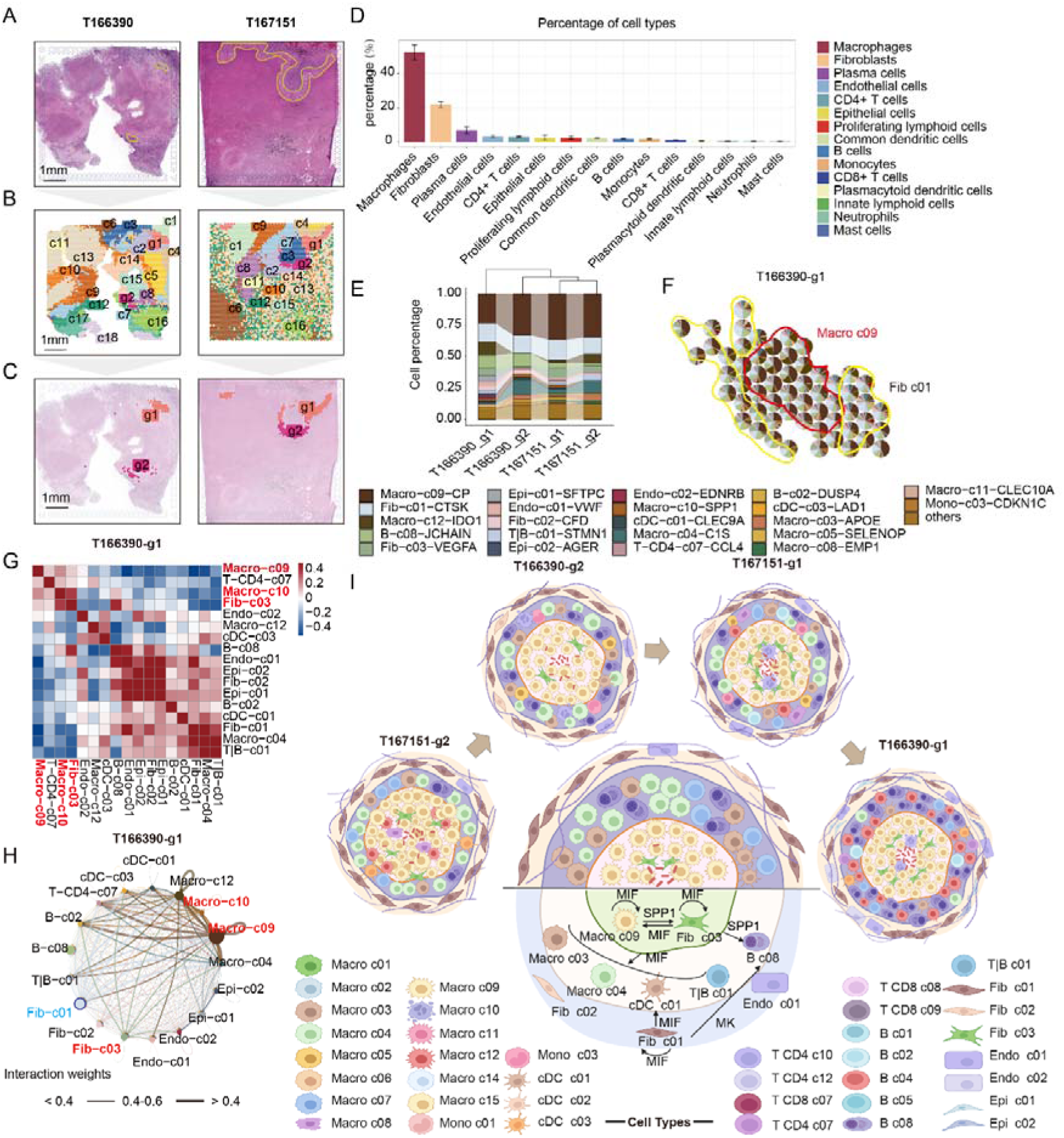
Cellular composition and spatial architectures of pulmonary granulomas. **(A)** Pulmonary lesions stained with H&E showing granuloma histological and histopathological regions (encircled by yellow lines). **(B-C)** Spatial visualization with spots representing whole lesions and four granuloma regions, color-coded by clusters. Spot positions match tissue locations. **(D)** Bar plot displaying the average proportion of each cell type within the four pulmonary granulomas. **(E)** Bar chart showing the cellular subpopulation distribution across four pulmonary granulomas. **(F)** Scatter pie plot showing the spatial cellular subpopulation composition of T166390-g1 granuloma. Each spot is represented as a pie chart. The red and yellow lines encircle the core and peripheral regions, respectively, showing the main cellular subpopulations. **(G)** Heatmap displaying cellular subpopulation co-localizations using Pearson correlations. **(H)** Circle plot depicting putative ligand-receptor interactions among different cellular populations, with the edge width representing the communication strength. **(I)** Spatial cellular three-layer architecture of pulmonary granulomas showing primary cellular populations and intercellular signaling communication (Created with BioRender.com). The central model is a representation that integrates primary cellular subpopulations and intercellular signaling present in at least three granulomas. The figure only showing signaling pathways among core layer cells and signals emitted from core layer and peripheral layer cells to intermediate layer cells.

Employing Robust Cell Type Decomposition (RCTD), we identified the cell-type composition of each spot based on scRNA-seq data. Macrophages (∼52.36%), fibroblasts (∼21.76%), and plasma cells (∼6.78%) were the top three cell types (Fig. 4D and fig. S7A, B). Further analysis of granulomas revealed that Macro-c09 (∼32.08%) and Fib-c01 (∼14.62%) were the main subtypes (fig. S7C-E; table S3). Intriguingly, granulomatous regions contained fewer B-c08 cells, suggesting structural barriers against plasma cell infiltration.

Among the four identified granuloma clusters, Euclidean distance analysis revealed that three clusters share similar cellular compositions. However, one cluster, T166390-g1, displayed significant heterogeneity (ED: 0.18 ∼ 0.2). This cluster is characterized by a higher presence of immune-activated Macro-c12 cells, suggesting a dynamic immune response to *Mtb* stimuli (Fig. 4e) (*30*).

### Three layers of spatial cellular architecture in pulmonary granulomas

To investigate cellular spatial distribution within granulomas, we conducted co-localization and intercellular communication analyses (Fig. 4F-I and fig. S8A-C). The granuloma structure is distinctly organized into three layers: core, middle, and periphery (Fig. 4I).

Macro-c09 cells are predominantly located in the core layer, often co-localizing with Fib-c03 cells, while Fib-c01 cells primarily occupied peripheries (Fig. 4F-G and fig. S8A-C). The three cell clusters showed strong pro-inflammatory activity and emitted robust signals—MIF, SPP1, and MK — to recruit and activate diverse immune cells into the middle layer (Fig. 4H, I; table S4) (*31–34*).

Furthermore, co-localization analysis revealed granulomas’ heterogeneity (Fig. 4I). In T167151-g1 and T166390-g1, Macro-c09 and Fib-c03 cells co-localized with Macro-c10 in the core. In T167151-g2, Macro-c09 and Fib-c03 co-localized with both Macro-c12 and Macro-c08 in the core. The middle layer exhibited greater heterogeneity, characterized by the co-existence of various immune cells, including inactive macrophages (Macro-c03/c04/c05), active macrophages (Macro-c08/c11/c12), cDCs (c01/03), CD4+T cells, and B (c02/c08) cells.

To depict typical features of pulmonary granulomas, we constructed a new granuloma model using cell types found in over half of the four granulomas (Fig. 4I), shedding light on general spatial communication between the layers.

### Significant enrichment of immune and metabolic pathways in pulmonary granulomas

Using gene set variation analysis (GSVA), we identified key pathways in granulomas (fig. S8D). Five pro-inflammatory pathways (IL2_STAT5, IL6_JAK_STAT3 signaling, inflammatory response, IFN-γ response and TNF-α/NF-κB signaling) were significantly enriched in the granulomas, indicating their crucial role in the host’s defense against *Mtb* (*35, 36*). Conversely, T166390-g1/g2 showed the enrichment of some anti-inflammatory pathways, like TGF_BETA and OXIDATIVE_PHOSPHORYLATION (*37, 38*). This duality in pro- and anti-inflammatory responses reflects an immune balance within granulomas as prior research (*39, 40*).

Additionally, these granulomatous clusters owned some significantly enriched metabolic pathways (fatty acid metabolism, glycolysis, ROS, hypoxia, etc.), known to modulate immune cell behavior for favoring bacterial survival (*41–44*). We also detected elevated pathways linked to cellular growth/proliferation (MTORC1 signaling, p53 pathways, adipogenesis, etc.), and DNA damage in granulomas, possibly resulting from the granuloma’s dense structure and heightened metabolic activity (*41*).

### Three-layer spatial cellular architecture of lymphatic granulomas with strong heterogeneity

We then explored the spatial cellular organization of lymphatic granulomas. Three lymphatic lesion samples (T166353, T166682, and T166210) were clustered into 14, 11, and 19 clusters, respectively. Pathological evaluations pinpointed a distinct cluster in each sample that corresponded to a well-defined granuloma region (Fig. 5A-C).

**Fig. 5.**
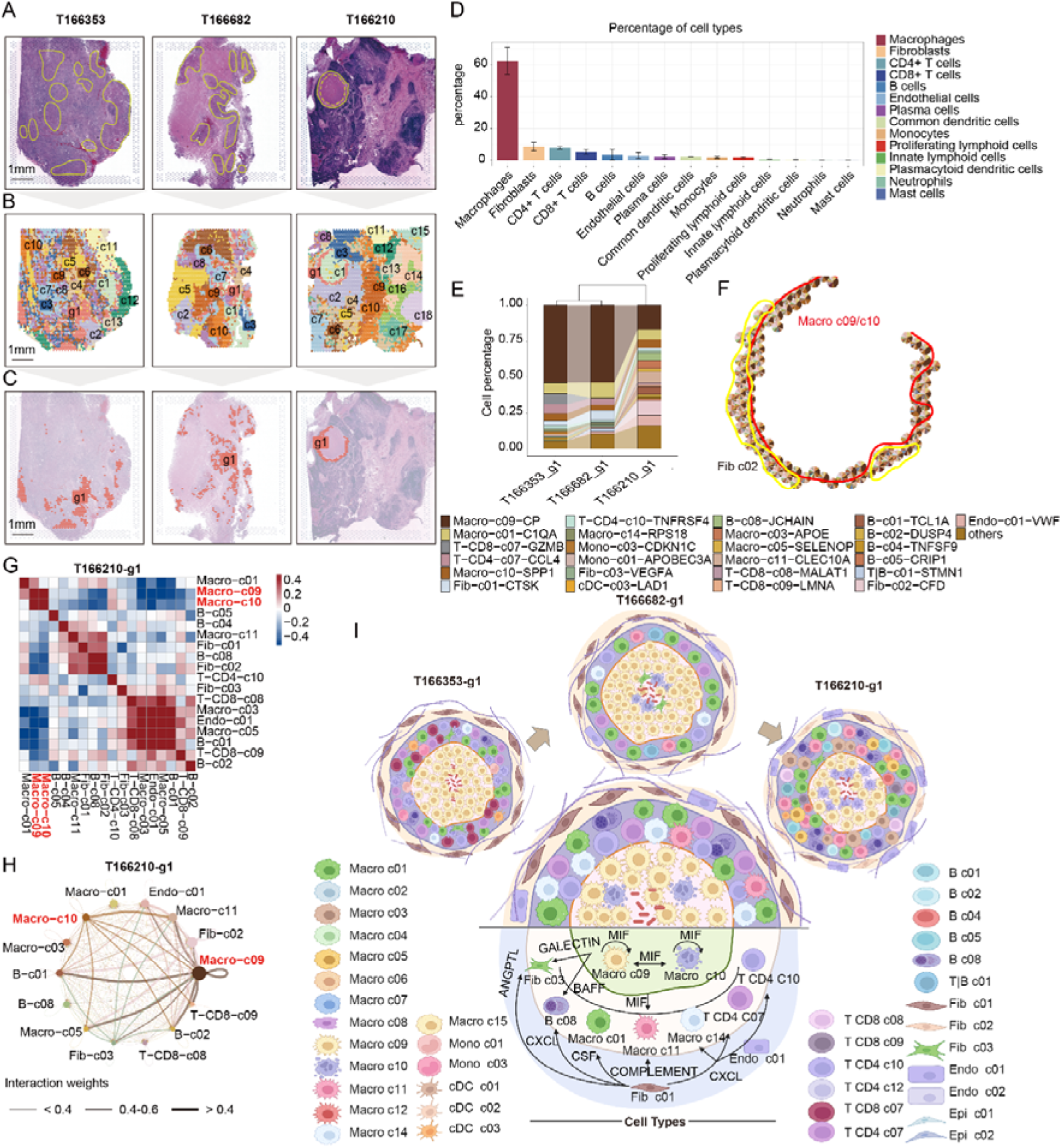
Cellular composition and spatial architectures of lymphatic granulomas. **(A)** Lymphatic lesions stained with H&E showing granuloma histological and histopathological regions. **(B-C)** Spatial visualization with spots representing whole lesions and three granuloma regions. **(D)** Bar plot displaying the average proportion of each cell type within the three lymphatic granulomas. **(E)** Bar chart showing the cellular subpopulation distribution across three lymphatic granulomas. **(F)** Scatter pie plot showing the spatial cellular subpopulation composition of T166210-g1 granuloma. **(G)** Heatmap displaying cellular subpopulation co-localizations. **(H)** Circle plot depicting putative ligand-receptor interactions among different cellular populations, with the edge width representing the communication strength. **(I)** Spatial cellular three-layer architecture of lymphatic granulomas showing primary cellular populations and intercellular signaling communication (Created with BioRender.com). The central model is a representation that integrates primary cellular subpopulations and intercellular signaling present in at least two granulomas. The figure only showing signaling pathways among core layer cells and signals emitted from core layer and peripheral layer cells to intermediate layer cells.

RCTD analysis indicated macrophages as the predominant cell type (over 50%), followed by fibroblast (∼8.67%). Lymphatic samples contained higher proportions of CD4+T cells (∼7.82%) and CD8+T cells (∼5.27%) than pulmonary samples, likely due to lymphatic tissue’s unique immune environment (Fig. 5D and fig. S9A-C). In lymphatic granulomas, Macro-c09 (∼41.55%) was predominant, paralleling observations in pulmonary granulomas (fig. S9D-G; table S5). The T166210-g1 granuloma diverged significantly in cell composition from the other two granulomas (ED: ∼0.4), with more Fib-c02 cells (∼10.5%), possibly linked to its caseous phenotype (Fig. 5E).

We further delineated the spatial cellular organization of lymphatic granulomas (Fig. 5F-I). Similar to pulmonary granulomas, Macro-c09 cells predominantly occupied the core, whereas Fib-c01 cells mainly occupied the periphery (fig. S10A-C). Core and peripheral cells emitted strong molecular signals (MIF, COMPLEMENT, CXCL, etc.) to recruit diverse immune cells to the middle layer (*45–47*) (Table S6). Compared to pulmonary granulomas, the lymphatic granulomas’ middle layer had a greater density of lymphocytes and monocytes, aligning with lymphatic tissue’s inherent immune properties.

We also observed marked spatial heterogeneity in three lymphatic granuloma regions (Fig. 5E-I). Specifically, T166353-g1’s core only consisted of Macro-c09 cells; T166682-g1 had a mixture of Macro-c09, Macro-c10 and Fib-c03 in the core; T166210-g1 displayed a denser co-localization of Macro-c09 with Macro-c10 in its core. Regarding the periphery, T166353-g1 and T166682-g1 were dominated by Fib-c01, while T166210-g1 showed a higher proportion of Fib-c02. These variations in the cellular composition of the core and periphery suggest differing states of granuloma activity among the samples.

GSVA analysis showed that lymphatic granulomatous regions (fig. S10D) were characterized by enriched pathways associated with immune response, metabolism, and cell growth/proliferation, paralleling those found in pulmonary granulomas (fig. S8D). Collectively, we characterized the general spatial cellular architecture of lymphatic granulomas (Fig. 5I), and highlighted an increased T and B cell presence.

### Three-layer spatial cellular structure of skeletal granulomas

Three skeletal lesion samples (T166457, T166846 and T167126) were clustered into 15, 18 and 9 distinct clusters, respectively (Fig. 6A-C). Five granulomatous regions were identified through expert pathological assessments: T166457 and T166846 each contained one, while T167126 had three (g1-g3). RCTD analysis showed macrophages (∼67.7%), fibroblasts (∼12.0%), and CD4+T cells (∼9.6%) as the dominant cell-types in the three skeletal lesion samples (Fig. 6D; fig. S11A-C). Further analysis indicated that Macro-c02 (∼34.5%) and Macro-c10 (∼11.7%) were the primary cell clusters of the skeletal granulomas (fig. S11D-G; table S7), differing from the pulmonary and lymphatic granulomas (Fig. 4E, 5E). Notably, an ascending trend of Macro-c10 cells from clusters g1 to g3 in T167126 suggests progressive disease severity (Fig. 6E).

**Fig. 6.**
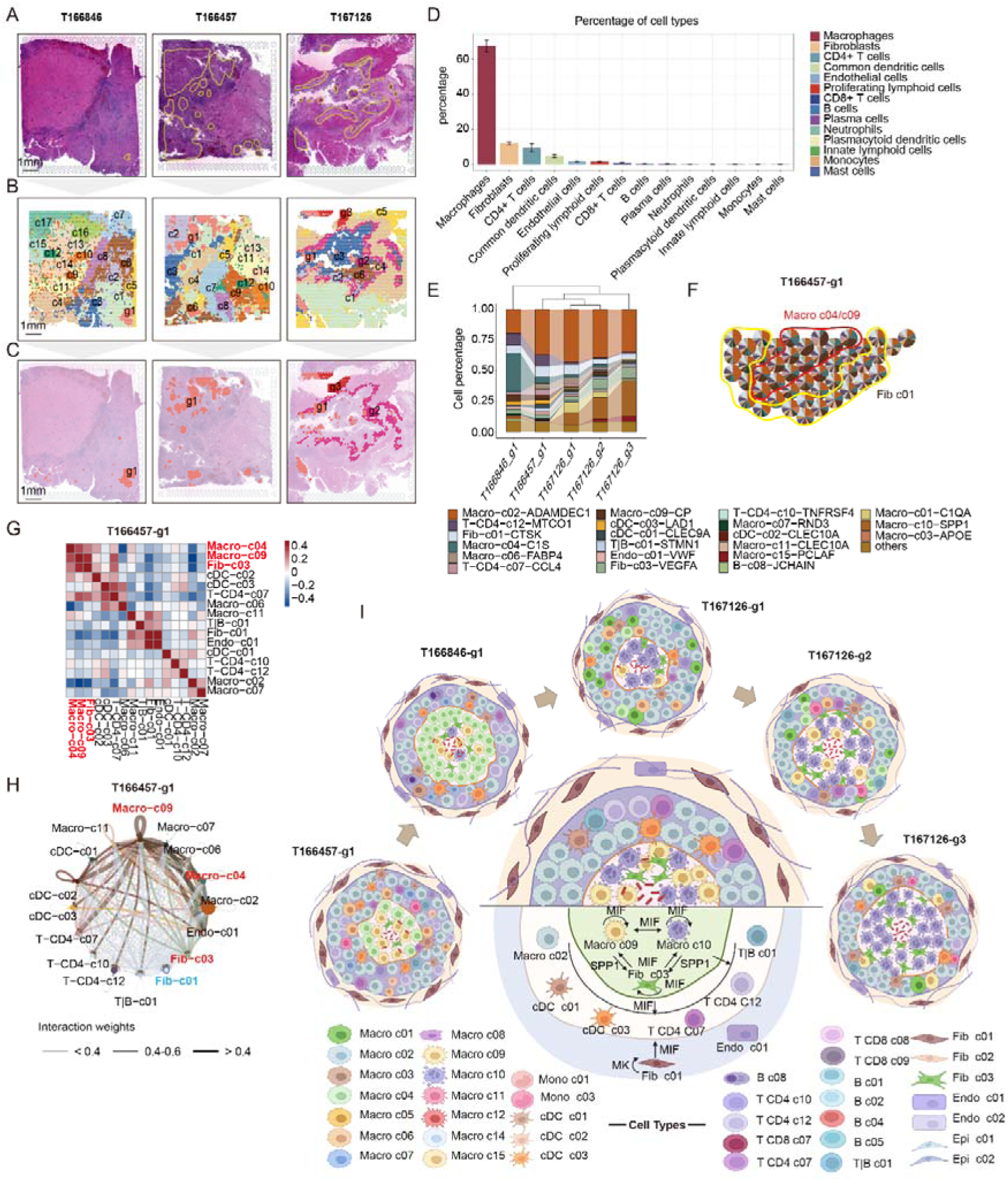
Cellular composition and spatial architectures of skeletal granulomas. **(A)** Skeletal lesions stained with H&E showing granuloma histological and histopathological regions. **(B-C)** Spatial visualization with spots representing whole lesions and five granuloma regions. **(D)** Bar plot displaying the average proportion of each cell type within the five skeletal granulomas. **(E)** Bar chart showing the cellular subpopulation distribution across five skeletal granulomas. **(F)** Scatter pie plot showing the spatial cellular subpopulation composition of T166457-g1 granuloma. **(G)** Heatmap displaying cellular subpopulation co-localizations. **(H)** Circle plot depicting putative ligand-receptor interactions among different cellular populations, with the edge width representing the communication strength. **(I)** Spatial cellular three-layer architecture of skeletal granulomas showing primary cellular populations and intercellular signaling communication (Created with BioRender.com). The central model is a representation that integrates primary cellular subpopulations and intercellular signaling present in at least two granulomas. The figure only showing signaling pathways among core layer cells and signals emitted from core layer and peripheral layer cells to intermediate layer cells.

Co-localization and intercellular communication analyses revealed a three-layer spatial structure for skeletal granulomas (Fig. 6F-I; fig. S12A-C). Similar to pulmonary and lymphatic granulomas, Macro-c09 and Fib-c03 primarily occupied the core, and Fib-c01 was enriched in the periphery. Differently, skeletal granulomas featured Macro-c02 in their middle layer, emitting strong GALECTIN signals to diverse immune cells (e.g., Macro-c09/c10, cDC-c01/c03, T-CD4-c07/c12, etc.), known to induce apoptosis and prompt inflammatory cytokine release (*48*) (table S8).

The heterogeneity was also observed, reflecting immune dynamics within skeletal granulomas during TB response (Fig. 6I). T166457-g1’s core contains both Macro-c09 and Fib-c03 alongside the less-inflammatory Macro-c04. T166846-g1 appears an increased presence of Macro-c10 in its core. Notably, T167126-g1 to g3 displayed an increasing proportion of Macro-c10, a trend that corresponds with pathological observations of increasing necrosis (fig. S6). These findings support our hypothesis that Macro-c10 represents a cell cluster transitioning towards cell death from Macro-c09.

GSVA results for skeletal granulomas mirrored pulmonary and lymphatic findings (such as immune, metabolic, and cellular growth/proliferation), highlighting common pathways essential for balanced granuloma inflammation across tissues (fig. S12D). Particularly, T167126 displayed decreased inflammatory but rising hypoxia signals from g1 to g3, indicating escalating severity and a transition from a non-necrotizing (g1) to necrotizing state (g2 and g3).

In conclusion, skeletal granulomas share certain spatial cellular architecture and functional characteristics with pulmonary and lymphatic granulomas, reflecting the consistency of TB granulomas. However, compared to pulmonary and lymphatic granulomas, the heightened presence of Macro-c10 denotes their distinct severity of skeletal ones, highlighting the individuality of TB granulomas across different tissues.

### Significantly up- and down-regulated genes in granulomas

To explore genetic marker genes indicative of TB granulomas, we analyzed differentially expressed genes (DEGs) between granulomatous and non-granulomatous regions within each sample (Data file S1). By assessing the occurrence of each DEG across 12 granulomas, we identified several marker genes (Table 1).

**Table 1.** Significantly up-/down-regulated marker genes within over 30% of the granulomas.

|  |  | Lung granulomas |  |  |  | Lymph granulomas |  |  | Bone granulomas |  |  |  |  | Frequency<br>of occurrence<br>in granulomas |
| --- | --- | --- | --- | --- | --- | --- | --- | --- | --- | --- | --- | --- | --- | --- |
| Markers |  | T167151<br>g2 | T166390<br>g2 | T167151<br>g1 | T166390<br>g1 | T166353<br>g1 | T166682<br>g1 | T166210<br>g1 | T166846<br>g1 | T166457<br>g1 | T167126<br>g1 | T167126<br>g2 | T167126<br>g3 |  |
| Up<br>regulated | FBP1 | - | 1.51 | 1.54 | 1.82 | 1.02 | - | 3.67 | 3.02 | 2.53 | - | 0.59 | - | 8 |
|  | SPP1 | - | - | 1.81 | - | 1.60 | - | 2.54 | 3.03 | 2.31 | - | - | 1.00 | 7 |
|  | G0S2 | 2.18 | - | 1.12 | - | 1.30 | - | - | - | 1.97 | - | 0.89 | - | 5 |
|  | CHI3L1 | - | 1.56 | - | - | - | - | 2.67 | 2.19 | - | 1.30 | - | 0.83 | 5 |
|  | MT1G | - | - | - | 1.74 | 1.32 | - | - | 3.13 | 2.22 | - | - | 0.98 | 5 |
|  | MMP9 | 3.44 | - | 1.57 | - | 1.04 | - | - | 2.74 | - | - | - | - | 4 |
|  | IDO1 | - | - | - | - | 1.01 | 0.55 | - | 1.93 | 2.20 | - | - | - | 4 |
| Down<br>regulated | IGKC | -0.77 | - | -1.60 | -0.94 | -1.91 | -1.16 | - | -2.15 | -3.29 | -2.34 | -1.77 | -1.61 | 10 |
|  | IGHG2 | -2.51 | - | - | -1.38 | -1.93 | -1.09 | - | -2.27 | -3.51 | -1.23 | -0.84 | -1.24 | 9 |
|  | IGHG1 | -3.13 | - | -0.56 | -1.02 | -1.67 | - | - | - | -3.34 | -1.39 | -1.25 | -1.11 | 8 |
|  | IGHG3 | -2.44 | - | -0.98 | - | -1.93 | -1.14 | - | - | -3.25 | - | - | - | 5 |
|  | IGLC1 | -0.78 | - | -0.49 | - | -1.78 | -1.07 | - | - | -3.28 | - | - | - | 5 |
|  | JCHAIN | - | -0.69 | -0.77 | - | -1.53 | -1.05 | - | - | -3.39 | - | - | - | 5 |
|  | IGHA1 | - | - | -2.00 | - | -1.81 | -1.26 | - | - | -3.65 | - | - | - | 4 |
|  | IGKV4-1 | - | - | -1.25 | - | -1.61 | -1.30 | - | -2.14 | - | - | - | - | 4 |
|  | HBA2 | - | - | - | - | - | -1.19 | - | - | - | -2.31 | -2.63 | -2.00 | 4 |

Seven genes (FBP1, SPP1, G0S2, CHI3L1, MT1G, MMP9, and IDO1) were significantly upregulated in over 30% of the granulomas, suggesting their crucial role in granuloma formation and the host’s immune response to *Mtb* invasion. Previous studies have reported the important roles of SPP1, MMP9 and IDO1 in TB granuloma formation (*49–52*). Newly discovered genes (FBP1, G0S2, CHI3L1, and MT1G) may contribute to granuloma formation through metabolic reprogramming and inflammatory processes, warranting further research (*53–56*).

Conversely, nine genes—IGKC, IGHG2, IGHG1, IGHG3, IGLC1, JCHAIN, IGHA1, IGHKV4-1, and HBA2—were significantly downregulated in over 30% of the granulomas. Notably, eight of them are related to immunoglobulins, indicating a significant reduction in antibody presence within granulomas. This aligns with prior observations of fewer plasma cells in granulomatous regions (fig. S7), highlighting the challenges the innate immune system faces against *Mtb*.

## Discussion

### Three-layer spatial structure of human TB granulomas

TB granulomas, the most defining morphological feature of TB, serve as the primary battleground between *Mtb* and its human host. Traditional histological studies have outlined a two-layered structure: a central core of macrophages surrounded by a ring of lymphocytes and fibroblasts (*57, 58*). However, a deeper understanding of their intricate architecture and cellular diversity is crucial.

Our scRNA-seq analysis identified nine key functional cell clusters characterized by robust immune activation and/or cell signaling (Fig. 2 and Fig. 3). Further, integrating ST-seq and scRNA-seq data, we proposed novel three-layer spatial cellular architectures for human TB granulomas (Fig. 4I, 5I, and 6I): the core layer is majorly occupied by Macro-c09 and Macro-c10, with occasional Fib-c03 cells; the peripheral layer is mainly Fib-c01, with some endothelial cells; and the intermediate layer displays a varied collection of immune cells, particularly macrophages and lymphocytes.

This research introduces two significant advancements in understanding human TB granulomas’ spatial structure. To our knowledge, we combined scRNA-seq with ST-seq to unveil an intricate three-layer structure of human TB granulomas for the first time. Secondly, we provide a comprehensive view of the cellular types, functions, intercellular communications, and significantly upregulated/downregulated genes within these layers, offering in-depth insights into cell clusters and even single-cells and molecules in the unique immune-microenvironment.

### Formation and dynamic evolution mechanisms of three-layer TB granulomas

Our integrative analysis of cellular distribution, composition, immune functions, and signaling interactions shed light on the formation and evolutionary dynamics of TB granulomas. Initially, we revealed key roles of Macro-c09 and Macro-c10 in the granuloma initiation, serving as primary host cells for *Mtb*, given their dominant core presence, substantial outgoing signaling, enhanced proinflammatory responses, and elevated SPP1 expression—a macrophage marker signaling of *Mtb* infection (*33, 49*). Additionally, a subset of Fib-c03 cells within the core is also immune-activated by robust SPP1 signals from adjacent Macro-c09 and Macro-c10. These three cell clusters constitute the granuloma core, collectively emitting strong signals to recruit and activate other cell clusters.

Further, Fib-c01, predominantly located in the periphery, is activated by receiving intense SPP1 signaling from Macro-c09 and Macro-c10, exhibiting strong signal emission and proinflammatory activity. Collectively, these core and periphery cells emit strong signals (like MIF, SPP1, CXCL and CCL) to recruit diverse immune cells for immunoinfiltration, including myeloid cells and lymphocytes, forming the granuloma’s intermediate layer (Fig. 4I, 5I, and 6I). Considerable heterogeneity in cell clusters within this layer was observed across granulomas, suggesting a degree of non-specific recruitment.

Our findings offer several novel insights into human granuloma development: (1) We provide a deepened understanding of granuloma formation and dynamic evolution at cell-cluster and molecular levels, focusing on characteristic cellular types, recruitment and activation mechanisms, and signaling molecules. (2) Our research pioneers in highlighting the pivotal roles of fibroblasts (Fib-c01 and Fib-c03) in granuloma development, demonstrating their enhanced proinflammatory responses and immune signal transduction for recruiting other immune cells. (3) Unlike prior fragmented insights (*9, 33, 58*), our study offers a comprehensive map of the signaling molecules in cellular communication during granuloma formation, presenting potential targets for future host-directed therapies.

### Heterogeneity among human TB granulomas

Our study reveals the heterogeneity among human TB granulomas. Previous studies on *Mtb*-infected non-human primates and patients also identified heterogeneity within TB granulomas, including differences in histopathological features (*12, 59, 60*), metabolic profiles (*61, 62*), inflammatory signals (*7*), and pathogen burdens (*9*).

Notably, we found that an increased presence of Macro c10 and Fib-c03 in the granuloma core correlates with granuloma necrosis progression, suggesting their potential as markers for disease advancement. Single-cell analysis also suggests that Macro-c10 might be a dysfunctional variant of Macro-c09 (Fig. 2F). Notably, the highest Macro-c10 presence was detected in skeletal granulomas’ core (Fig. 6I), consistent with clinical reports suggesting skeletal TB’s often more severe nature compared to its pulmonary and lymphatic counterparts, typically manifesting with tissue necrosis and abscesses.

Additionally, we identified tissue-related differences in cellular composition and spatial structure across lymphoid, lung, and skeletal granulomas (Fig. 4-6). Lymphoid granulomas showed higher levels of immune cell infiltration, whereas lung and skeletal ones had more immune-activated fibroblasts in their cores, reflecting the lymphatic system’s naturally higher immune cell density and lower fibroblast count. Casanova et al.’s recent work on human sarcoidosis lymph node granulomas aligns with our observations, highlighting the impact of location on immune response (*63*). Overall, our data suggest that inter-individual variations in granulomas are often more pronounced than tissue-related differences.

Our contributions to TB granuloma heterogeneity include: (1) introducing a novel comparative analysis at high resolution (single-cell levels), exploring spatial cellular composition differences within and between tissue types, advancing beyond previous studies that focused solely on comparisons within the same tissue; (2) conducting a comprehensive heterogeneity analysis based on full transcriptomic genes, providing a broader view than prior studies limited to select marker genes.

While our study advances the understanding of human tuberculous granulomas’ structure and function, its limitations should be acknowledged. The granuloma samples were collected from patients who required surgery due to drug-resistant tuberculosis. The limited number of these surgical specimens underscores the need for more comprehensive future studies to enhance representativeness. Additionally, our ability to conduct further experimental validations is constrained by the lack of necessary licensing for experiments on cells or animals infected with *Mtb*, as well as the limited availability of samples.

## Methods

### Study design

The primary objective of this study is to dissect the intricate cellular composition, spatial architecture, intercellular communications, and immune functions of human TB granulomas. The study participants were recruited from Beijing Chest Hospital, Capital Medical University between March and September 2022 (table S1). Granulomatous tissue specimens were selected from pulmonary, lymphatic and skeletal tuberculosis patients based on two main criteria: 1) The need for surgical removal of the granulomas from patients resistant to conventional drug treatments; 2) Histological confirmation of chronic granulomas with necrosis via H&E staining; and 3) Patients were excluded if they had liver or autoimmune diseases, were pregnant or lactating, or had a history of prolonged use of steroids or other immunosuppressive agents. Finally, 11 specimens (4 pulmonary, 5 lymphatic and 2 skeletal) were collected for scRNA-seq, and 8 specimens (4 pulmonary, 2 lymphatic and 2 skeletal) were collected for ST-seq. For collected samples, we further identified granuloma areas in each slice by locating regions rich in epithelioid and giant cells, and assessed granulomas’ number, size, location, and necrosis extent. By integrating scRNA-seq with spatial-seq, we identify nine key cell clusters and describe a consistent three-layer spatial cellular architecture, with similar cell composition, functions and intercellular communications, but also uncover heterogeneity likely linked to disease severity.

### Clinical specimens

Each resected tissue was divided into two equal parts: one part, measuring approximately 1cm x 1cm x 2cm, was used for single-cell transcriptomics; the other part, also measuring 1cm x 1cm x 2cm, was used for ST-seq. The latter samples were first fixed with 10% formaldehyde, subsequently dehydrated, and embedded with paraffin (FFPE samples) for downstream experiments (fig. S13).

### scRNA-seq library preparation and sequencing

To prepare each sample, approximately 1.0 – 1.5 g of tissue was minced into smaller pieces (∼0.5 mm^2^ per piece) using scissors. Excessive blood was removed from the homogenates by adding 35 ml of ice-cold PBS and gently shaking the mixture, followed by filtration through a 40 μm strainer to collect the tissue. The tissue was then treated with an enzyme mixture containing dispase (50 U/ml), collagenase (2 mg/ml), elastase (1 mg/ml), and DNase (30 μg/ml) for enzymatic digestion at 37 °C for 1 h with shaking. The reaction was terminated by adding 5 ml of PBS supplemented with 10% FCS. The dissociated cells were passed through a 70 μm strainer and centrifuged at 300 g for 5 min at 4 °C. The resulting cell pellet was resuspended in 3 ml of red blood cell lysis buffer and incubated at room temperature for 2 min to lyse remaining red blood cells. After incubation, 10 ml of PBS supplemented with 10% FCS was added to the suspension, followed by centrifugation at 300 g for 5 min at 4 °C.

Single-cell suspensions were subjected to single-cell sample processing and cDNA library preparation following the instructions provided by the manufacturer (10x Genomics). Samples with a viability above 90% were selected for scRNA-seq, using either the Chromium Single Cell 30 (v3) Reagent Kit or the Chromium Single Cell 59 Library and Chromium Single Cell V(D)J Amplification Kits, Human (10x Genomics), in accordance with the manufacturer’s standard protocols. The constructed 3’ expression, 5’ expression, or TCR/BCR V(D)J libraries were sequenced on a NovaSeq sequencing platform (Illumina) with a target of 50,000 read pairs per cell.

### scRNA-seq data processing

The scRNA-seq data from each of the 3’ expression libraries were processed using the count pipeline of the Cellranger toolkit (v6.0), which included alignment against the GRCh38 human reference. For the matched 5’ expression and TCR/BCR V(D)J libraries, the Cellranger multi-pipeline was employed to simultaneously analyze and aggregate the data. Cells with less than 1000 UMIs, less than 500 detected genes, or more than 10% mitochondrial genes were excluded using the Seurat package v4.0 (*64*). Scrublet (*65*) was used to identify potential cell doublets using the default parameters, except for an expected doublet rate of 0.1. Cells with UMI count above 50,000 or detected genes above 5,000, as well as putative doublets flagged by Scrublet, were filtered out. Notably, the CD3E+ myeloid cell cluster was identified as doublets and removed, as it expressed both myeloid gene signature (CST3) and T cell signature (CD3E).

### Dimension reduction, unsupervised clustering, cell-type determination and further analysis

The Seurat package was utilized to further process the filtered data. To eliminate batch effects, the Harmony algorithm was used to merge data from different samples based on the first 30 principal components (PCs). The merged dataset was subjected to UMAP dimensionality reduction and clustering with the 30 PCs and a resolution of 1.0. Cluster-specific marker genes with Benjamini-Hochberg (BH) adjusted P-values lower than 0.05 were identified, and cell types were manually annotated based on the cluster markers. A second-round sub-clustering was performed on myeloid cells, lymphoid cells, and stromal cells following the above-described strategy. The first 30 PCs and a resolution of 2.0 were employed for cell clustering and UMAP dimension reduction. The same method was used to identify marker genes, which were subsequently utilized to annotate and phenotype the cell sub-populations. Notably, previously published signatures (table S2) were also utilized for T cell phenotyping, whereas CellTypist was used for B cell phenotyping.

GSEA analyses were performed using the clusterProfiler package (*66*) (v3.14.3) with gene sets from the Molecular Signatures Database (MsigDB) Hallmark collection, GO knowledgebase, and KEGG database. Gene sets, terms or pathways were considered as statistically significant if their BH-adjusted P-values were lower than 0.05. RNA velocity analyses were performed based on spliced and unspliced reads for each individual cell using the scVelo pipeline with default parameters. Cell-cell communication analysis was performed using CellChat (*67*) (v0.5) with default parameters. Clonotype analysis of TCR and BCR was performed using the scirpy package in Python.

### ST-seq library preparation and sequencing

Following the guidelines of 10X Genomic Visium Spatial Gene Expression for FFPE, ST-seq libraries were generated for sequencing. For each sample, the FFPE tissue block was sectioned into 10 µm slices and placed onto the capture area of a Gene Expression slide (10X Genomics). The slides were then subjected to H&E staining and scanned for imaging. Next, ST-seq libraries were constructed using the Visium Spatial Gene Expression Reagent Kit. Finally, the libraries were sequenced on Illumina NovaSeq 6000 sequencing platform.

### Spatial sequencing analysis

To determine the location information of the capture area for each sample, Loupe Browser (v6.1.0) was used to manually calibrate the image of the capture area, and a .json file containing the location information was generated. The ST-seq data was processed using the SpaceRanger toolkit (v1.3.1), which included alignment against the GRCh38 human reference. A feature (gene expression)-barcode (spot) was generated for each library to describe the gene expression values of each spot. Subsequently, Seurat v4.1.3 was used to analyze the expression matrix of each sample separately. Normalization was performed using the SCTransform method, and UMAP dimensionality reduction was performed using the RunUMAP function with the first 30 PCs. For clustering of spots, the first 30 PCs and resolutions of 0.6 to 1.8 were used. To achieve a more precise delineation of the granulomatous areas, a second-round sub-clustering was performed on the granulomatous clusters identified in the initial round for T166390 (resolution=2) and T167126 (resolution=0.5) samples. The GSVA analysis for the unsupervised clusters was further performed based on the MsigDB.

### Spot-level cell type decomposition

To determine cell type composition in each spot of the samples, the Robust Cell Type Decomposition (RCTD) method (*68*) (spacexr v2.0.3) was utilized by referring to the cell types identified in the scRNA-seq analysis. Pulmonary-specific cell types from scRNA-seq data labeled ST-seq spots in pulmonary tissue samples, lymphatic-specific cell types for lymphatic tissue, and skeletal-specific cell types for skeletal tissue. Average gene expression profiles for each cell type in annotated scRNA-seq data were computed, and deconvolution and maximum likelihood estimation techniques were used to infer the proportion of cell subtypes in each spot. The standard RCTD pipeline was applied with default parameters, except for "CELL_MIN_INSTANCE=25" and "doublet_mode=’full’".

### Spatial co-localization of cell types

To identify spatially co-localizing cell types, the R function "rcorr" was used to compute the Pearson correlation coefficient matrix for cell types within individual spots. This analysis included only cell types representing over 1% of the granulomatous regions. Cell type pairs with a correlation coefficient exceeding 0.5 were considered spatially co-localized.

### Statistical analysis

All data analyses were performed under R version 4.1.2 and Python 3.10.8. The statistical methods are fully described in the corresponding sections in the Methods and briefly outlined in the figure legends. False discovery rate correction was applied to p-values for multiple hypothesis testing as specified. A significance level of adjusted p < 0.05 was considered statistically significant.

## Acknowledgments

We acknowledge all study participants and clinical research staffs who contributed to this work. We thank Xiaotong Wang, Liya Yue, and Tianyi Lu for technical assistance. We thank Shuangshuang Li, Chenyang Wang, and Menlu Zhang for critical feedback. We also thank all the patients who allowed us to use their materials and made this work possible.

## Authors’ contributions

X.Y., J.W., P.W., X.L, C.L. contributed equally to this work. F.C. and H.R.H. conceptualized this study. J.W., P.H.W., X.Q.L., and C.D.L. wrote the original draft of the manuscript. X.Y., J.W., P.H.W., X.Q.L, C.D.L., and Y.J.Y. contributed to bioinformatics analyses, experimentation, data generation and visualization the manuscript. X.Y., Y.J.D., X.Y.J., B.T., H.W., W.B.Z., H.T.N., S.H.X., C.L.J., and J.W. contributed to investigation. F.C., J.W., P.H.W., X.Q.L., C.D.L., and Y.J.Y. revised the manuscript. All authors read and approved the final manuscript.

## Ethics approval and consent to participate

The study received ethical approval from the Ethics Committee of the Beijing Institute of Genomics, Chinese Academy of Sciences (China National Center for Bioinformation) (Approved number: 2023H006), and from the Beijing Chest Hospital, Capital Medical University (China) (Approved number: NO.BJXK-2022-KY-44). All patients enrolled in this study signed written informed consents prior to participation. The samplings and corresponding experiments were conducted in accordance with the approved ethical and biosafety protocols.

## Consent for publication

Not applicable.

## Availability of data and materials

The scRNA-seq and ST-seq data generated and analyzed in this manuscript have been deposited in the Gene Expression Omnibus with accession number: GSE271346. Meanwhile, these data have already been uploaded and are available at the Genome Sequence Archive (*69*) in the National Genomics Data Center, China National Center for Bioinformation/Beijing Institute of Genomics, Chinese Academy of Sciences, under accession number GSA-Human: HRA005957. Code used for the analysis of the scRNA-seq data and ST-seq data is available at https://doi.org/10.5281/zenodo.11634482.

