## Supplementary Materials for "Single-Cell and Spatial Transcriptomic Analyses Deciphering the Three-Layer Architecture of Human Tuberculosis Granulomas"

### Supplemental Figures

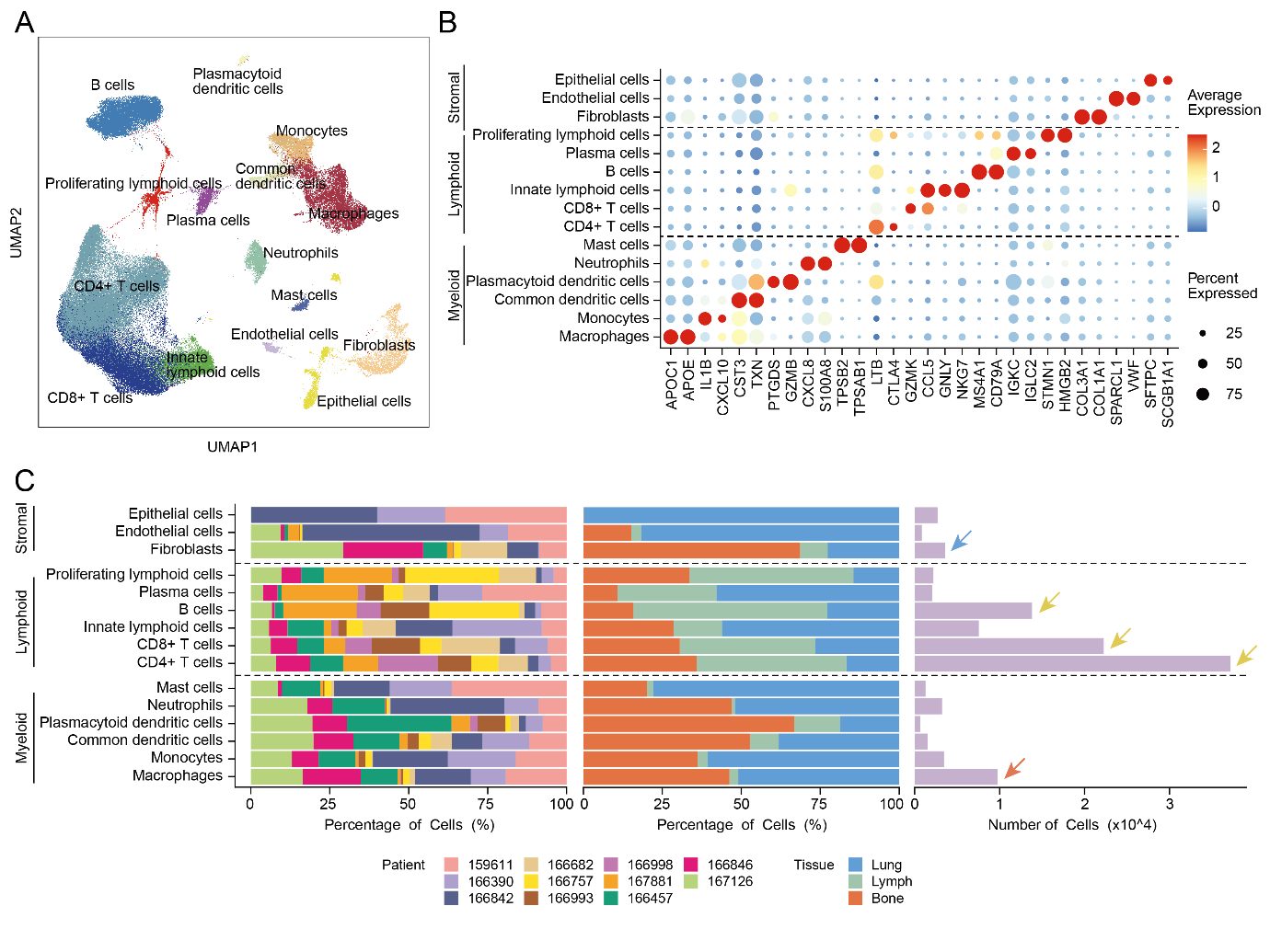

Fig. S1. scRNA-seq analysis of pulmonary, lymphatic and skeletal TB granuloma samples.

**A)** UMAP plot of 111,967 cells colored according to the identities of 15 major cell types. **B)** Dot plot illustrating expression levels of the genes defining the 15 major cell types. Color intensity corresponds to average gene expression, and dot size indicates the percentage of cells with expression in each of the 15 major cell types. **C)** Heterogeneity of the 15 major cell types displaying the cell percentage and number across different samples and tissues.

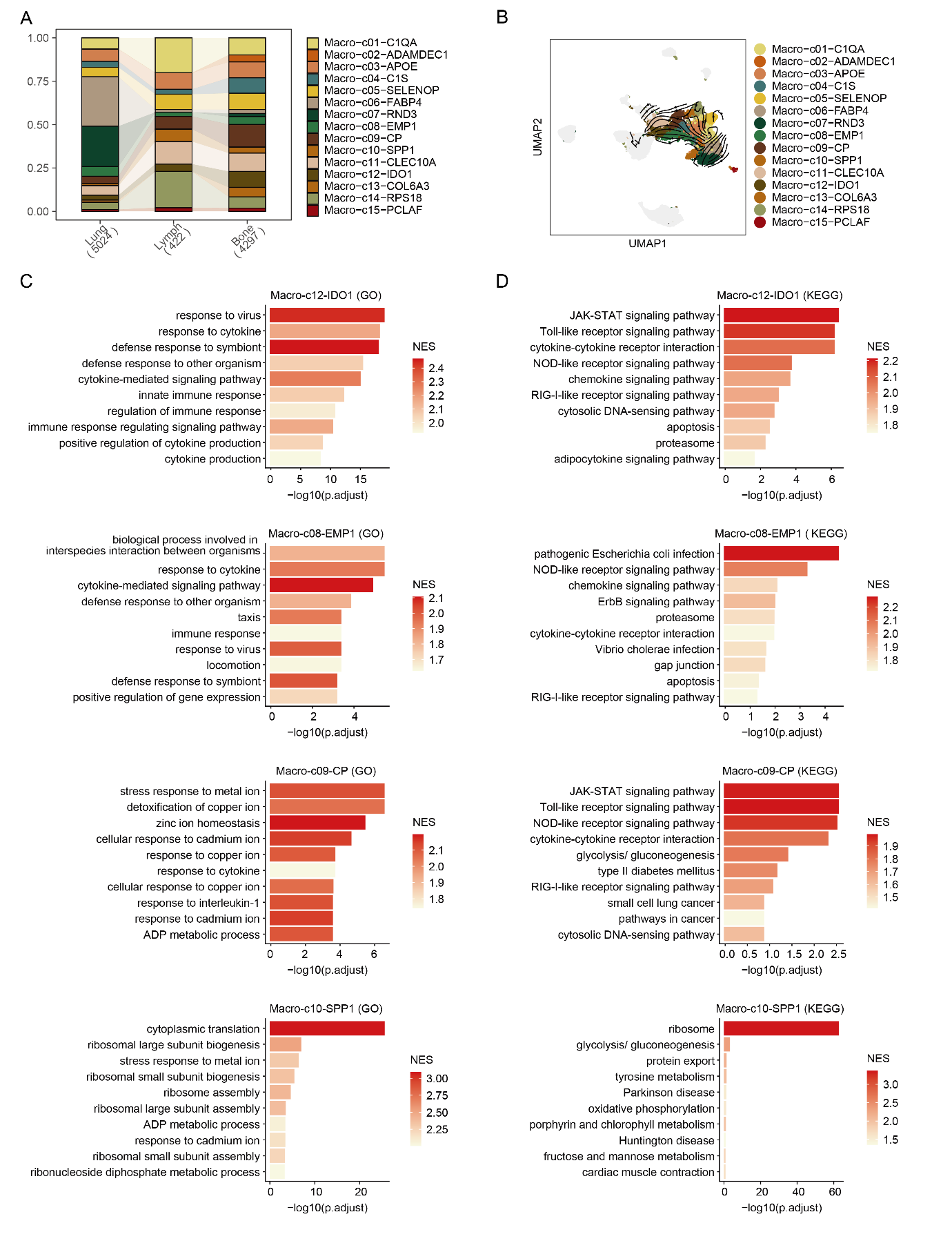

Fig. S2. Macrophage subpopulations in TB granuloma samples.

**A)** Bar plots showing macrophage subpopulation proportions across pulmonary, lymphatic, and skeletal TB granuloma samples. **B)** UMAP plot of macrophage subpopulation developmental transition revealed by RNA velocity. **C)** GSEA analyses for macrophage subpopulations Macro-c12, Macro-C08, Macro-c09 and Macro-c10 using GO gene sets. Color intensity corresponds to the level of NES. **D)** GSEA analyses for macrophage subpopulations Macro-c12, Macro-C08, Macro-c09 and Macro-c10 using KEGG gene sets. Color intensity corresponds to the level of NES.

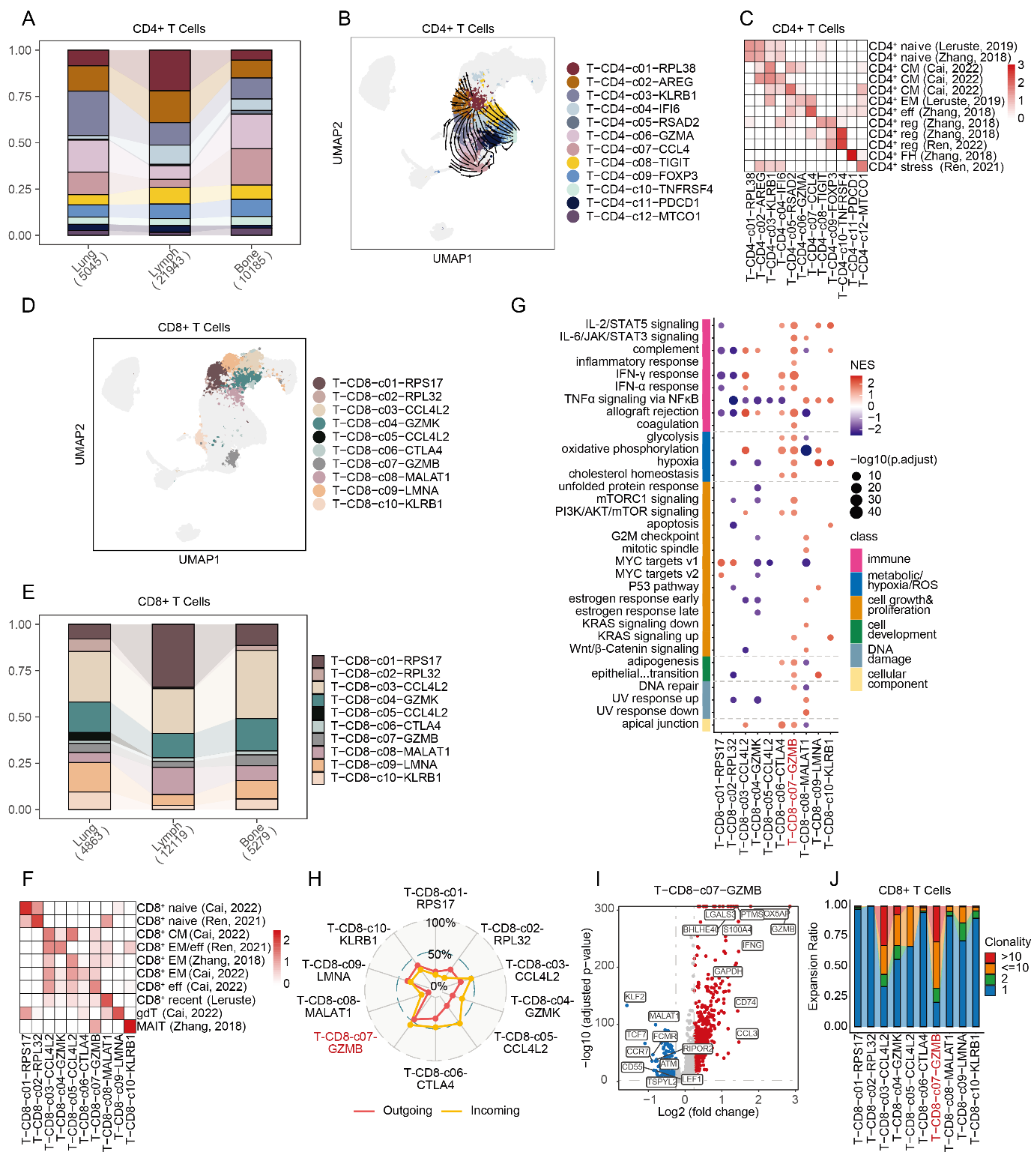

Fig. S3. CD4^+^ and CD8^+^ T cell subpopulations in TB granuloma samples.

**A)** Bar plots showing CD4^+^ T cell subpopulation proportions across pulmonary, lymphatic, and skeletal TB granuloma samples. **B)** UMAP plot of CD4^+^ T cell subpopulation developmental transition as revealed by RNA velocity. **C)** Heatmap showing enrichment of selected signatures or marker genes from published dataset for CD4^+^ T cell subpopulations. Color intensity corresponds to average enrichment level. **D)** UMAP plot of CD8^+^ T cells across all samples colored according to according to the identities of cell subpolulations. **E)** Bar plots showing CD8^+^ T cell subpopulation proportions across pulmonary, lymphatic, and skeletal TB granuloma samples. **F)** GSEA analyses for CD8^+^ T cell subpopulations using MSigDB hallmark gene sets. Color intensity represents NES, while dot size reflects statistical significance. **G)** Heatmap showing enrichment of selected signatures or marker genes from published dataset for CD8^+^ T cell subpopulations. Color intensity corresponds to average enrichment level. **H)** Radar plot showing intercellular interaction of CD8^+^ T cell subpopulations categorized as outgoing and incoming signaling. **I)** Volcano plots showing DEGs in T-CD8-c07 compared to other CD8^+^ T cells. Red and blue dots denote up-regulated and down-regulated DEGs, respectively. **J)** TCR clonality in CD8^+^ T cell subpopulations.

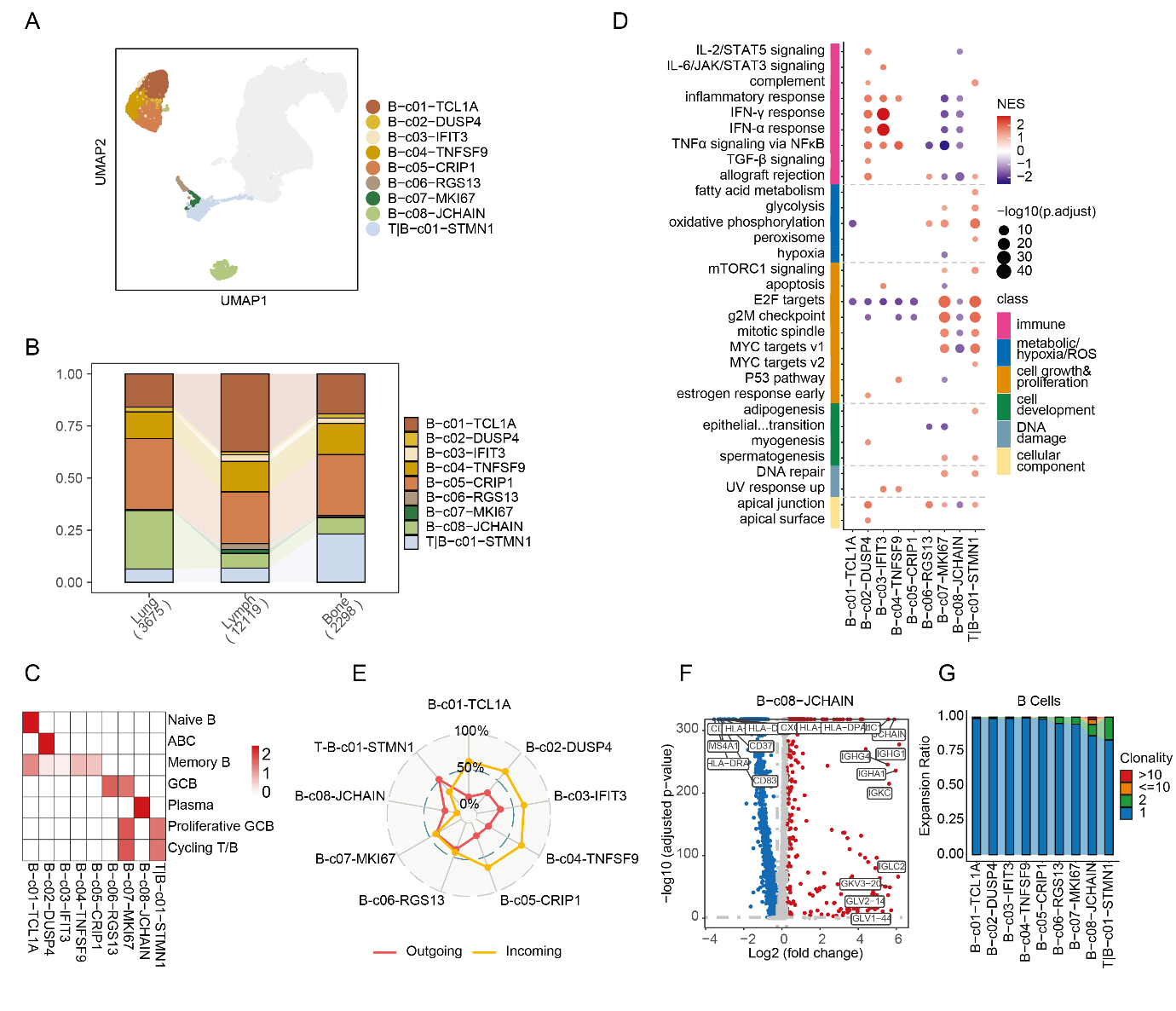

Fig. S4. B cell subpopulations in TB granulomas.

**A)** UMAP plot of B cells across all samples colored according to according to the identities of cell subpopulations. **B)** Bar plots showing B cell subpopulation proportions across pulmonary, lymphatic, and skeletal TB granuloma samples. **C)** GSEA analyses for B cell subpopulations using MSigDB hallmark gene sets. Color intensity represents NES, while dot size reflects statistical significance. **D)** Heatmap showing enrichment of selected signatures or marker genes from published dataset for B cell subpopulations. Color intensity corresponds to average enrichment level. **E)** Radar plot showing intercellular interaction of B cell subpopulations categorized as outgoing and incoming signaling. **F)** Volcano plots showing DEGs in B-c09 compared to other B cells. Red and blue dots denote up-regulated and down-regulated DEGs, respectively. **G)** BCR clonality in B cell subpopulations.

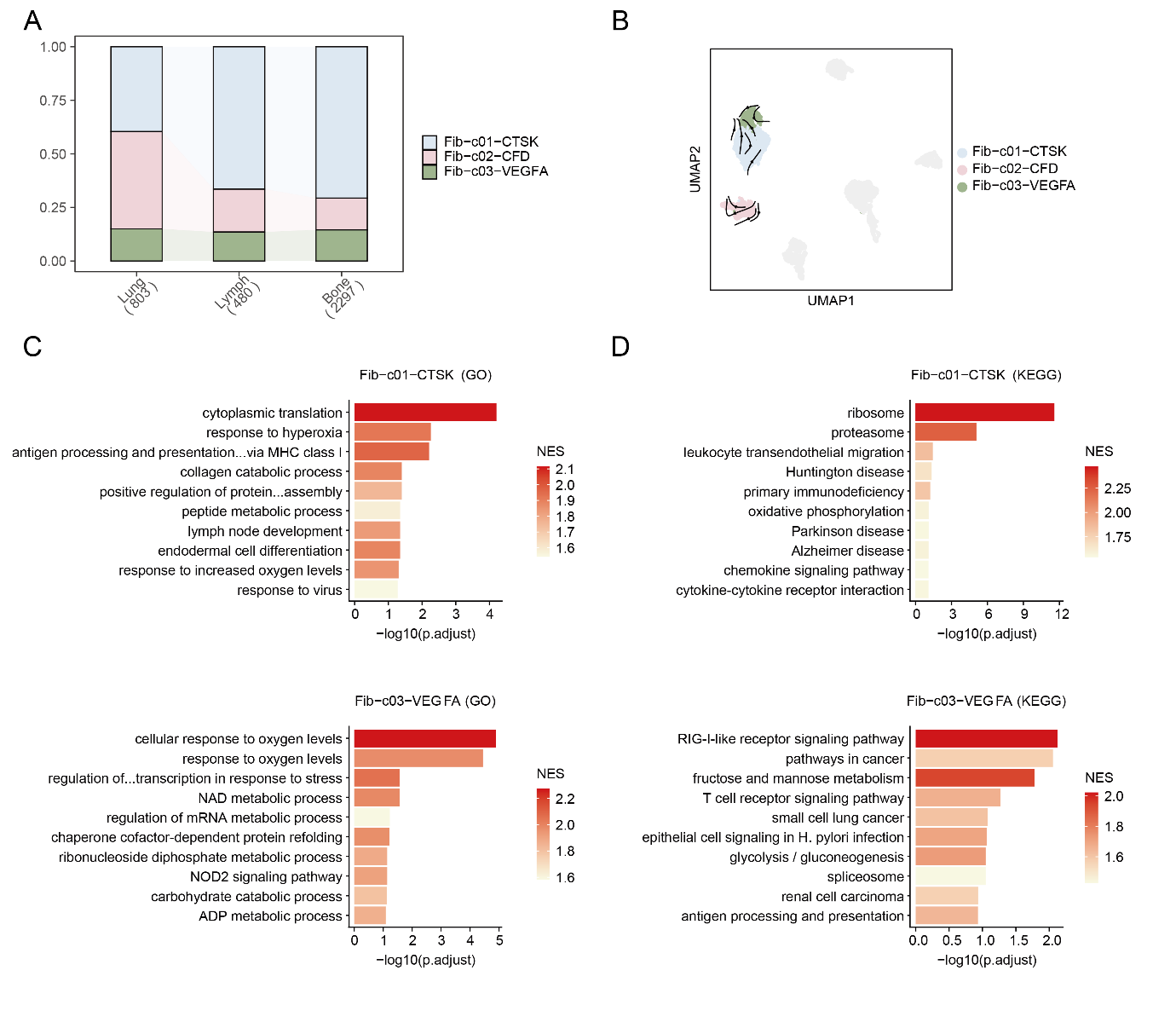

Fig. S5. Fibroblast subpopulations in TB granulomas.

**A)** Bar plots showing fibroblast subpopulation proportions across pulmonary, lymphatic, and skeletal TB granuloma samples. **B)** UMAP plot of fibroblast subpopulation developmental transition as revealed by RNA velocity. **C)** GSEA analyses for fibroblast subpopulations Fib-c01 and Fib-c03 using GO gene sets. Color intensity corresponds to the level of NES. **D)** GSEA analyses for fibroblast subpopulations Fib-c01 and Fib-c03 using KEGG gene sets. Color intensity corresponds to the level of NES.

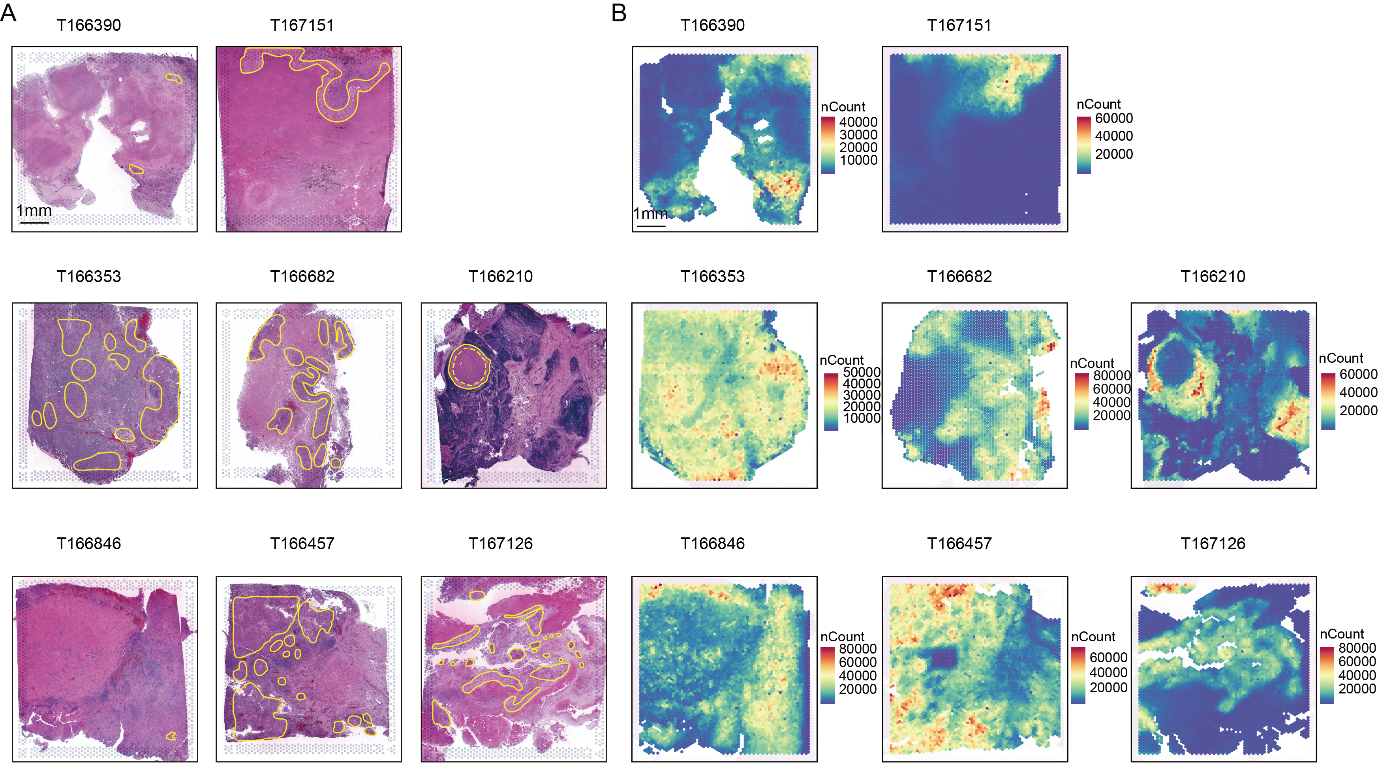

Fig. S6. Spatially resolved transcriptomic data statistics.

**A)** H&E-stained images of pulmonary, lymphatic, and skeletal TB lesions. Scale bar is 1 mm. **B)** Total counts of UMIs detected per spot.

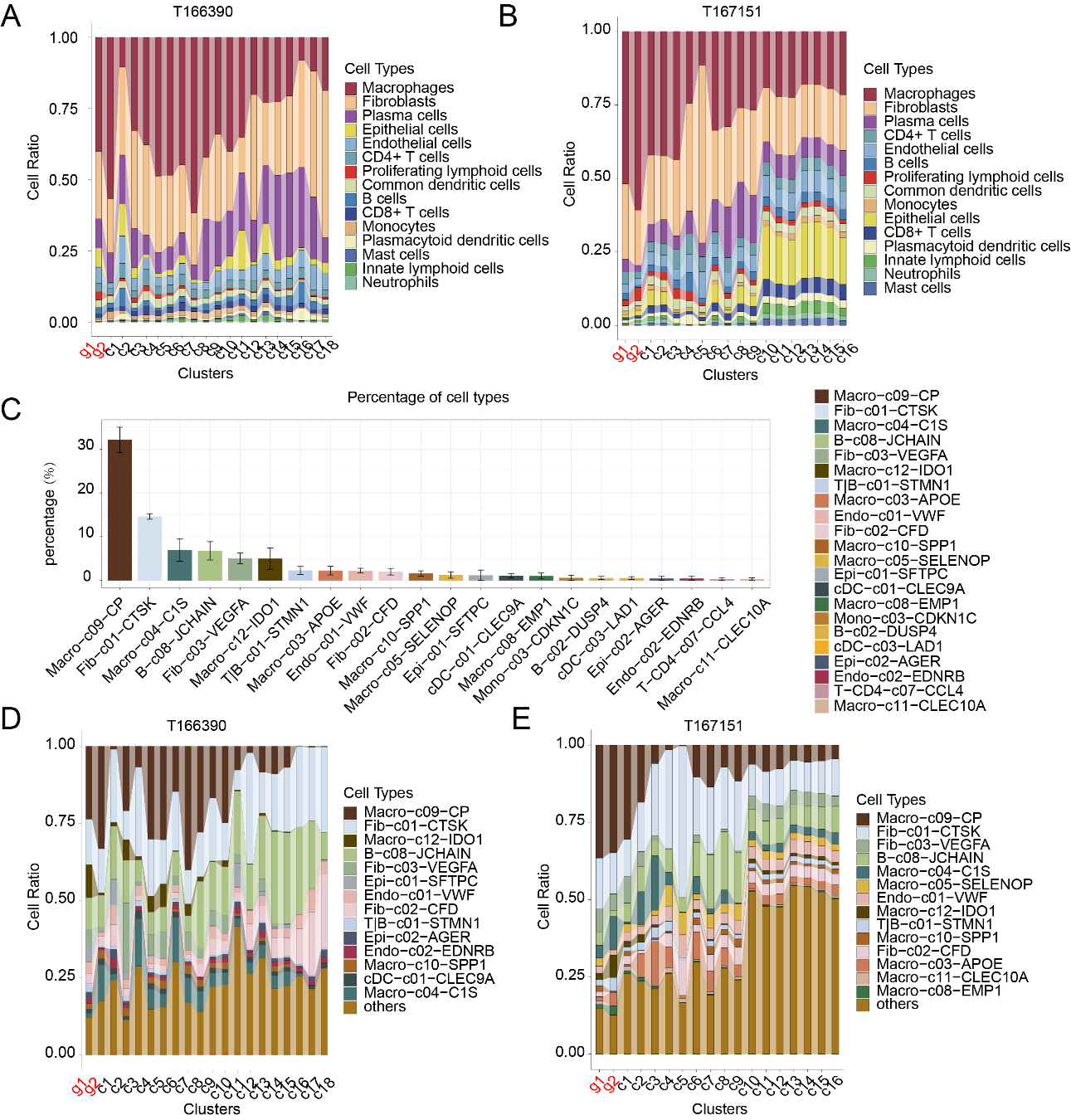

Fig. S7. Cellular composition of pulmonary granulomas.

**A and B)** Bar chart depicting the 15 major cell types distribution across different clusters of T166390 **(A)** and T167151 **(B)**, with the granulomatous regions grouped into clusters g1-g2. **C)** Bar plot displaying the average proportion of each cellular subpopulation within the four pulmonary granulomas. Only cellular subpopulation with a prediction accounting for over 1% in each spot were included. **D and E)** Bar chart displaying the top 14 cellular subpopulation distribution across different clusters of T166390 **(D)** and T167151 **(E).**

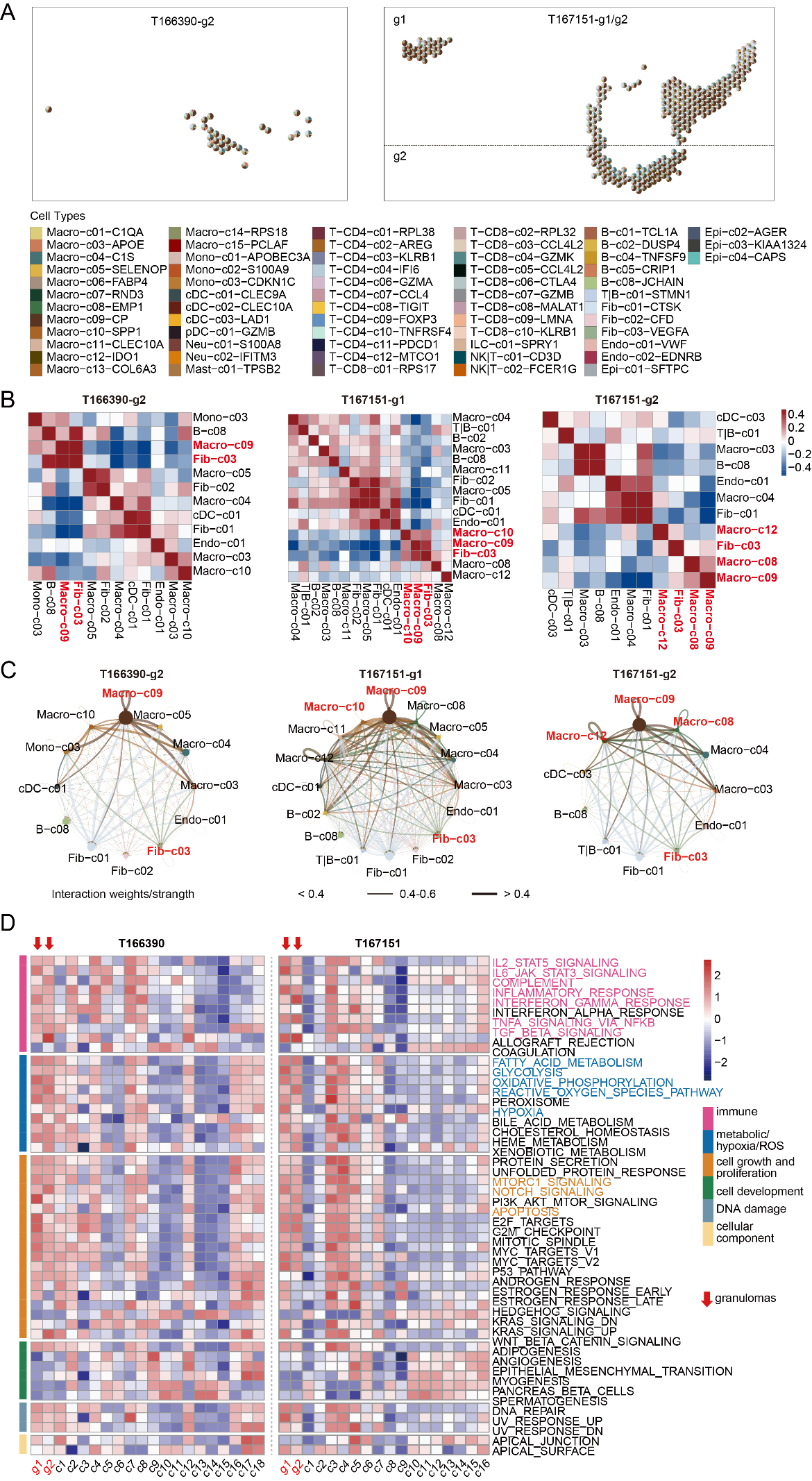

Fig. S8. Spatial architectures and functional profiling of granulomas in T166390 and T167151 samples.

**A)** Scatter pie plot showing the spatial cellular subpopulation composition of the T166390-g2 (left) and T167151-g1/g2 (right). Each spot is represented as a pie chart. **B)** Heatmap displaying cellular subpopulation co-localizations. **C)** Circle plots depicting putative ligand-receptor interactions among different cellular populations, with the edge width representing the communication strength. **D)** GSVA analyses for clusters of T166390 and T167151 samples using MSigDB hallmark gene sets.

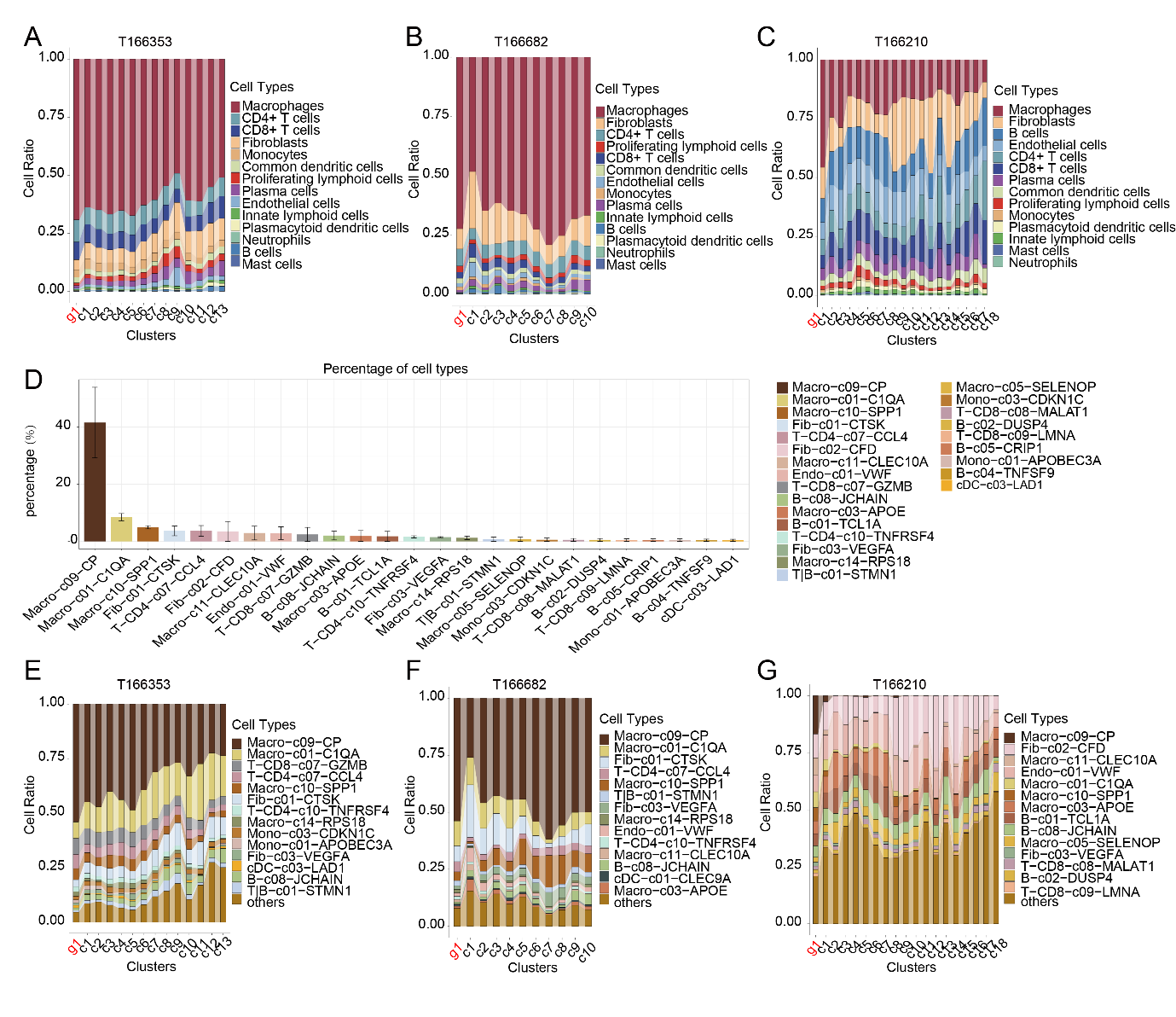

Fig. S9. **Cellular composition of lymphatic granulomas.**

**A, B and C)** Bar chart depicting the 15 major cell types distribution across different clusters of T166353 **(A)**, T166682 **(B)** and T166210 **(C)**. **D)** Bar plot displaying the average proportion of each cellular subpopulation within the three lymphatic granulomas. Only cellular subpopulation with a prediction accounting for over 1% in each spot were included. **E, F and G)** Bar chart displaying the top 14 cellular subpopulation distribution across different clusters of T166353 **(E)**, T166682 **(F)** and T167210 **(G).**

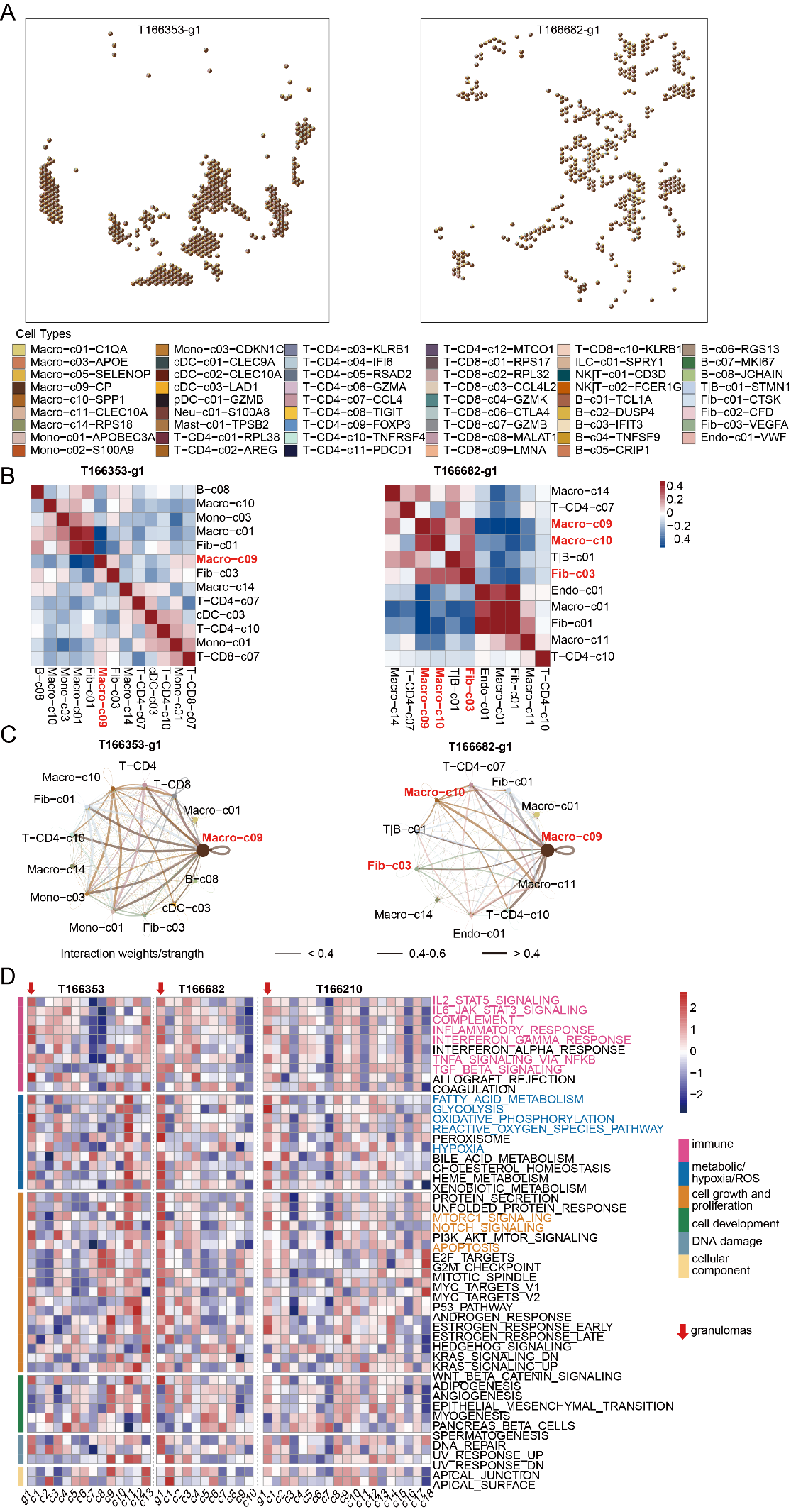

Fig. S10. **Spatial architectures and functional profiling of granulomas in T166353 and T166682 samples.**

**A)** Scatter pie plot showing the spatial cellular subpopulation composition of the T166353-g1 (left) and T166682-g1 (right). Each spot is represented as a pie chart. **B)** Heatmap displaying cellular subpopulation co-localizations. **C)** Circle plots depicting putative ligand-receptor interactions among different cellular populations, with the edge width representing the communication strength. **D)** GSVA analyses for clusters of T166353 and T166682 samples using MSigDB hallmark gene sets.

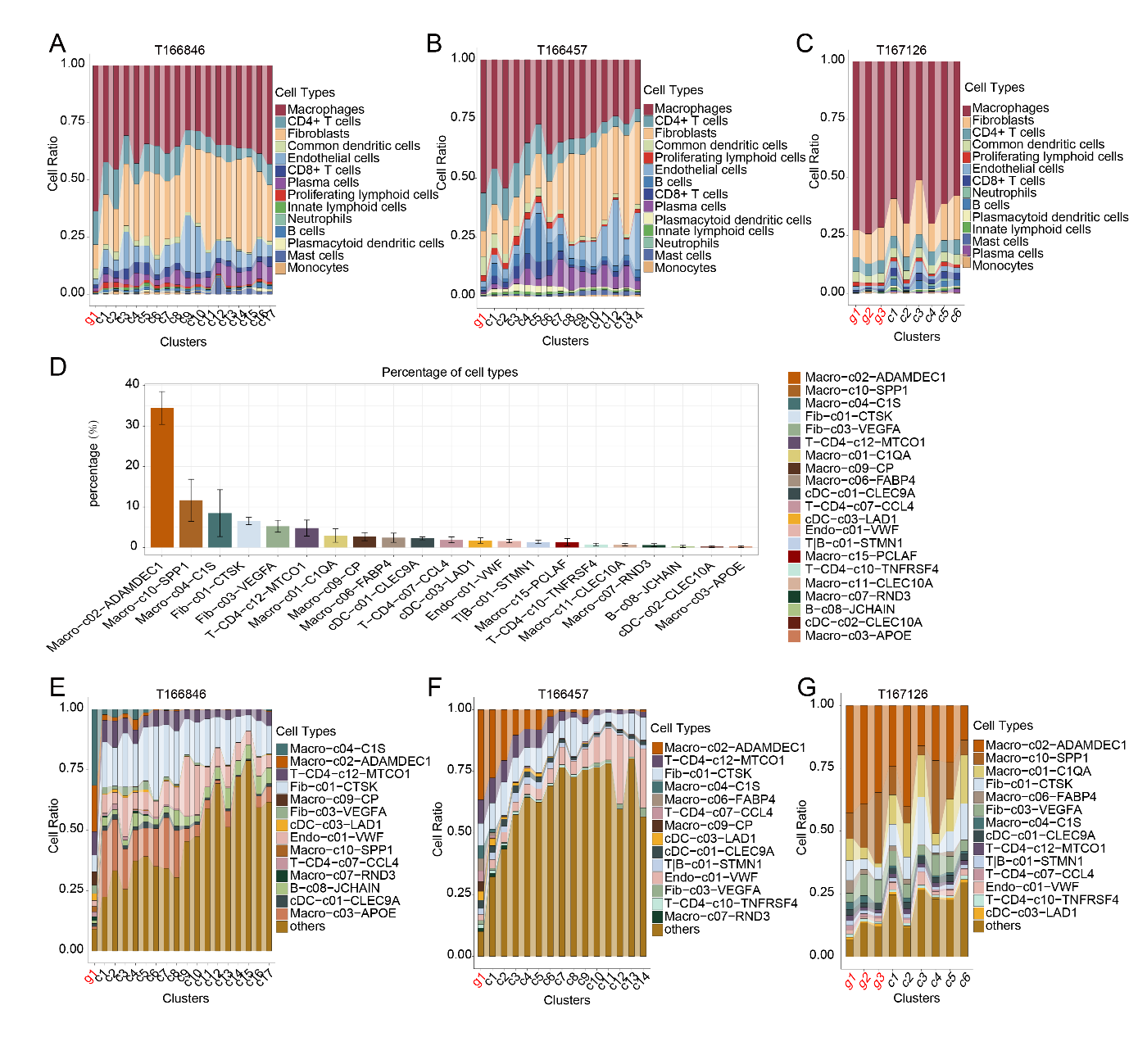

Fig. S11 **Cellular composition of skeletal granulomas.**

**A, B and C)** Bar chart depicting the 15 major cell types distribution across different clusters of T166846 **(A)**, T166457 **(B)** and T167126 **(C)**. **D)** Bar plot displaying the average proportion of each cellular subpopulation within the five skeletal granulomas. Only cellular subpopulation with a prediction accounting for over 1% in each spot were included. **E, F and G)** Bar chart displaying the top 14 cellular subpopulation distribution across different clusters of T166846 **(E)**, T166457 **(F)** and T167126 **(G).**

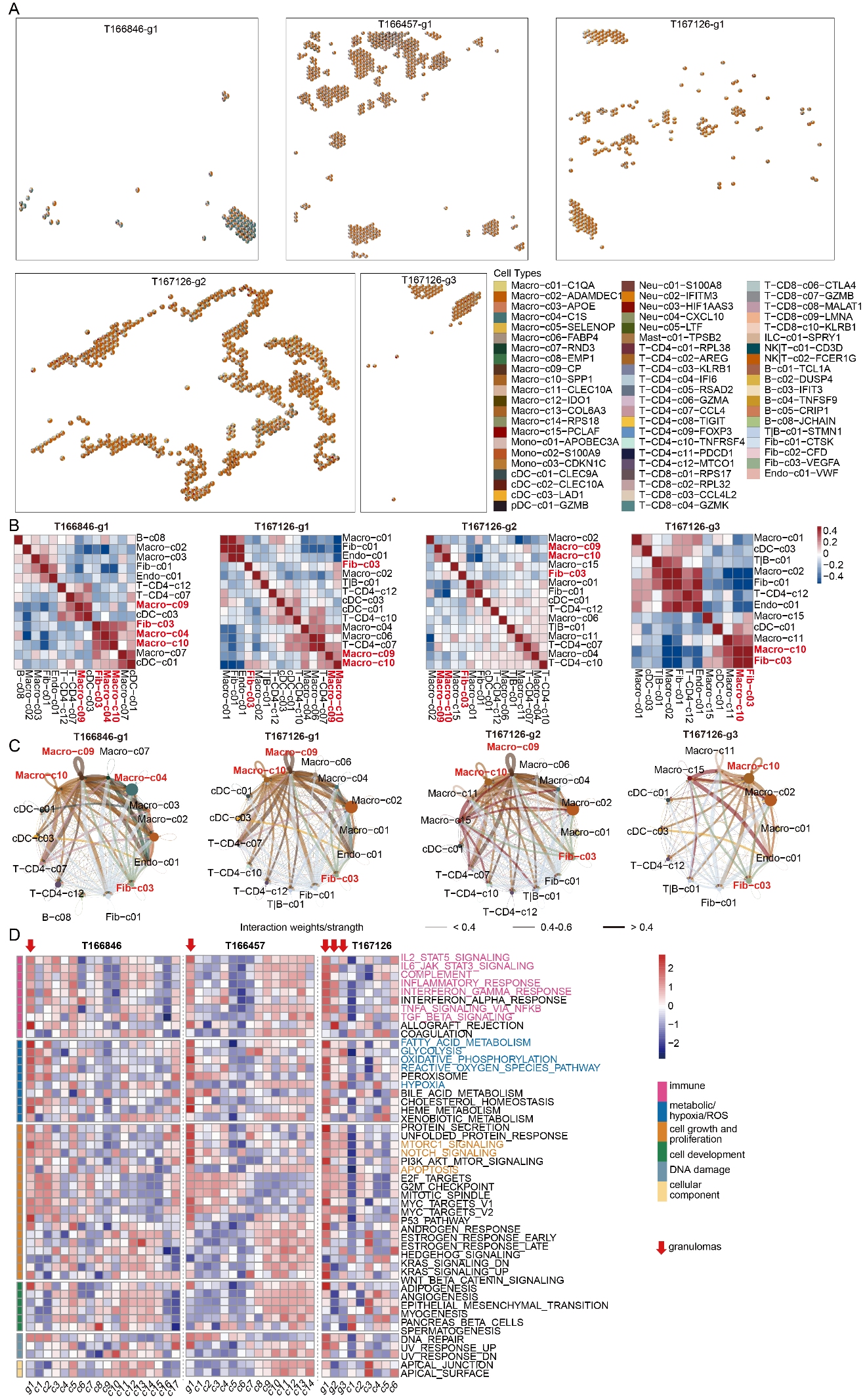

Fig. S12. **Spatial architectures and functional profiling of granulomas in T166846, T166457 and T167126 samples.**

**A)** Scatter pie plot showing the spatial cellular subpopulation composition of the T166846-g1, T166457-g1 and T167126-g1/g2/g3. Each spot is represented as a pie chart. **B)** Heatmap displaying cellular subpopulation co-localizations. **C)** Circle plots depicting putative ligand-receptor interactions among different cellular populations, with the edge width representing the communication strength. **D)** GSVA analyses for clusters of T166846, T166457 and T167126 samples using MSigDB hallmark gene sets.

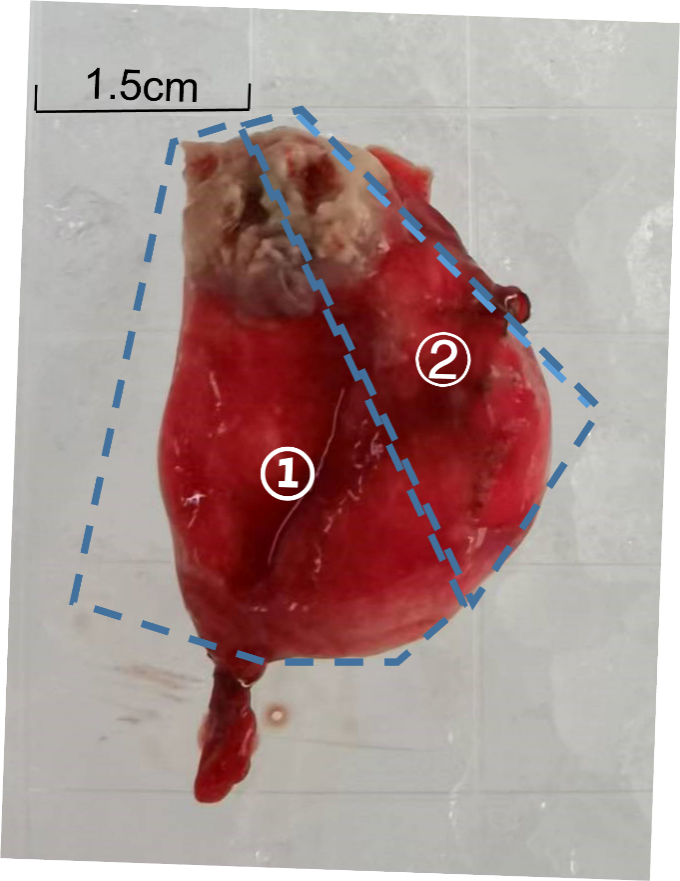

Fig. S13. Schematic of tissue dissection for multi-omics (sample ID 166390-lung).

Section ① was used for single-cell transcriptomics, measuring 1cm x 1cm x 2cm. Section ② was utilized for formalin-fixed paraffin-embedded (FFPE) procedures for pathological examinations and spatial transcriptome analysis, also measuring 1cm x 1cm x 2cm. Other samples were processed in the same manner as T166390.

### Supplemental Tables

Table S1. Basic information and sequencing details of single-cell and spatial transcriptomic samples.

| **Patient** | **Sex** | **Age** | **Tissue** | **Timing of sample collection** | **Library** | **Estimated number of cells** | **Mean  reads  per  cell** | **Median UMI per cell** | **Median genes  per cell** | **Number of spots under tissue** | **Fraction reads in spots under tissue** | **Mean reads per spot** | **Median UMI counts per spot** | **Median genes per spot** |
| --- | --- | --- | --- | --- | --- | --- | --- | --- | --- | --- | --- | --- | --- | --- |
| **159611** | Male | 20 | Lung | 06/30/22 | 3’ Expression | 14072 | 42,922 | 4,176 | 1528 |  |  |  |  |  |
|  |  |  |  |  | 5’ Expression | 11,248 | 20,171 | 4,728 | 1,819 |  |  |  |  |  |
| **166210** | Male | 21 | Lymph | 03/25/22 | Spatial |  |  |  |  | 4,705 | 98.50% | 40,110 | 4,256 | 2,342 |
| **166353** | Female | 29 | Lymph | 04/01/22 | Spatial |  |  |  |  | 4,264 | 96.70% | 43,650 | 22,008 | 5,460 |
| **166390** | Male | 30 | Lung | 04/02/22 | 3’ Expression | 9,166 | 68,211 | 4,779 | 1,804 |  |  |  |  |  |
|  |  |  |  |  | Spatial |  |  |  |  | 3,973 | 87.10% | 48,057 | 5,518 | 1,977 |
| **166457** | Male | 19 | Bone | 04/11/22 | 3’ Expression | 10,113 | 58,537 | 5,517 | 1,891 |  |  |  |  |  |
|  |  |  |  |  | Spatial |  |  |  |  | 4,558 | 95.80% | 48,243 | 24,654 | 5,954 |
| **166682** | Female | 48 | Lymph | 04/21/22 | 3’ Expression | 7,925 | 66,437 | 5,266 | 1,958 |  |  |  |  |  |
|  |  |  |  |  | Spatial |  |  |  |  | 3147 | 0.849 | 67,278 | 21056 | 5504 |
| **166757** | Female | 31 | Lymph | 04/19/22 | 3’ Expression | 9,709 | 57,845 | 4,468 | 1,629 |  |  |  |  |  |
| **166842** | Female | 49 | Lung | 05/16/22 | 3’ Expression | 13,569 | 45,871 | 2,463 | 1,172 |  |  |  |  |  |
| **166846** | Male | 57 | Bone | 05/09/22 | 3’ Expression | 12,284 | 50,089 | 6,259 | 2,142 |  |  |  |  |  |
|  |  |  |  | 04/26/22 | Spatial |  |  |  |  | 4,819 | 98.60% | 35,454 | 17,176 | 5,059 |
| **166993** | Female | 22 | Lymph | 04/27/22 | 3’ Expression | 11,072 | 47,194 | 6,232 | 1,914 |  |  |  |  |  |
| **166998** | Female | 27 | Lymph | 04/27/22 | 3’ Expression | 14,846 | 34,025 | 4,580 | 1,541 |  |  |  |  |  |
| **167126** | Female | 61 | Bone | 05/09/22 | 3’ Expression | 27,012 | 20,060 | 3,022 | 1,361 |  |  |  |  |  |
|  |  |  |  |  | Spatial |  |  |  |  | 4,248 | 87.50% | 40,608 | 6,767 | 2,782 |
| **167151** | Male | 53 | Lung | 05/13/22 | Spatial |  |  |  |  | 4,980 | 99.90% | 37,915 | 3,762 | 989 |
| **167881** | Female | 51 | Lymph | 10/06/22 | 5’ Expression | 14,629 | 18,572 | 4,790 | 1,840 |  |  |  |  |  |

Table S2. Reference gene signature list for CD4+ T cell, CD8+ T cell and B cells.

| **Cell Subpopulation** | **Correlated Publicatuion** | **Correlated Cell Subpopulation** | **Description** |
| --- | --- | --- | --- |
| T−CD4−c01−RPL38 | Leruste, 2019 | X1−CD4 | CD4+ naive |
| T−CD4−c02−AREG | Zhang, 2018 | CD4−C01 | CD4+ naive |
| T−CD4−c03−KLRB1 | Cai, 2022 | CD4−C06 | CD4+ CM |
| T−CD4−c04−IFI6 | Cai, 2022 | CD4−C03 | CD4+ CM |
| T−CD4−c05−RSAD2 | Cai, 2022 | CD4−C07 | CD4+ CM |
| T−CD4−c06−GZMA | Leruste, 2019 | X4−CD4 | CD4+ EM |
| T−CD4−c07−CCL4 | Zhang, 2018 | CD4−C09 | CD4+ eff |
| T−CD4−c08−TIGIT | Zhang, 2018 | CD4−C10 | CD4+ reg |
| T−CD4−c09−FOXP3 | Zhang, 2018 | CD4−C12 | CD4+ reg |
| T−CD4−c10−TNFRSF4 | Ren, 2022 | CD4−C11 | CD4+ reg |
| T−CD4−c11−PDCD1 | Zhang, 2018 | CD4−C11 | CD4+ FH |
| T−CD4−c12−MTCO1 | Ren, 2021 | T−CD4−c05 | CD4+ stress |
| T−CD8−c01−RPS17 | Cai, 2022 | CD8−C01 | CD8+ naive |
| T−CD8−c02−RPL32 | Ren, 2021 | T−CD8−c01 | CD8+ naive |
| T−CD8−c03−CCL4L2 | Cai, 2022 | CD8−C03 | CD8+ CM |
| T−CD8−c04−GZMK | Ren, 2021 | T−CD8−c06 | CD8+ EM/eff |
| T−CD8−c05−CCL4L2 | Zhang, 2018 | CD8−C04 | CD8+ EM |
| T−CD8−c06−CTLA4 | Cai, 2022 | CD8-C05 | CD8+ EM |
| T−CD8−c07−GZMB | Cai, 2022 | CD8-C07 | CD8+ eff |
| T−CD8−c07−GZMB | Leruste, 2019 | X20-CD8 | CD8+ recently activated |
| T−CD8−c09−LMNA | Cai, 2022 | gdT−02 | gdT |
| T−CD8−c10−KLRB1 | Zhang, 2018 | CD8−C08 | MAIT |

Table S3. The cell type composition of granulomas in two pulmonary tissue samples.

| Cell subpopulation | Rough classes | Pulmonary granulomas | | | | Cell subpopulation average |
| --- | --- | --- | --- | --- | --- | --- |
|  |  | T166390_g1 | T166390_g2 | T167151_g1 | T167151_g2 |  |
| Macro-c09-CP | Macropahge | 0.237 | 0.332 | 0.365 | 0.348 | 0.3208 |
| Fib-c01-CTSK | Fibroblast | 0.147 | 0.142 | 0.163 | 0.133 | 0.1462 |
| Macro-c04-C1S | Macropahge | 0.014 | 0.118 | 0.040 | 0.106 | 0.0695 |
| B-c08-JCHAIN | Plasma cell | 0.103 | 0.104 | 0.044 | 0.020 | 0.0678 |
| Macro-c12-IDO1 | Macropahge | 0.107 | 0.009 | 0.021 | 0.072 | 0.0520 |
| Fib-c03-VEGFA | Fibroblast | 0.059 | 0.015 | 0.075 | 0.053 | 0.0507 |
| T\|B-c01-STMN1 | T\|B cells | 0.028 | 0.004 | 0.020 | 0.044 | 0.0241 |
| Macro-c03-APOE | Macropahge | 0.000 | 0.035 | 0.013 | 0.043 | 0.0228 |
| Endo-c01-VWF | Endothelial cells | 0.038 | 0.013 | 0.023 | 0.017 | 0.0226 |
| Fib-c02-CFD | Fibroblast | 0.031 | 0.033 | 0.017 | 0.002 | 0.0207 |
| Macro-c10-SPP1 | Macropahge | 0.018 | 0.028 | 0.019 | 0.003 | 0.0170 |
| Macro-c05-SELENOP | Macropahge | 0.008 | 0.024 | 0.027 | 0.002 | 0.0150 |
| cDC-c01-CLEC9A | common dendritic cell | 0.014 | 0.019 | 0.010 | 0.005 | 0.0123 |
| Epi-c01-SFTPC | Epithelial cell | 0.047 | 0.000 | 0.000 | 0.001 | 0.0123 |
| Macro-c08-EMP1 | Macropahge | 0.003 | 0.002 | 0.012 | 0.029 | 0.0116 |
| Endo-c02-EDNRB | Endothelial cells | 0.019 | 0.006 | 0.008 | 0.006 | 0.0097 |
| Mono-c03-CDKN1C | Monocyte | 0.006 | 0.024 | 0.004 | 0.002 | 0.0090 |
| B-c02-DUSP4 | B cell | 0.013 | 0.007 | 0.010 | 0.004 | 0.0086 |
| T-CD4-c07-CCL4 | CD4+T cell | 0.013 | 0.006 | 0.008 | 0.004 | 0.0076 |
| cDC-c03-LAD1 | common dendritic cell | 0.011 | 0.003 | 0.005 | 0.011 | 0.0075 |
| Epi-c02-AGER | Epithelial cell | 0.019 | 0.001 | 0.004 | 0.006 | 0.0074 |
| pDC-c01-GZMB | Plasmacytoid dendritic cells | 0.003 | 0.005 | 0.009 | 0.006 | 0.0058 |
| T-CD4-c12-MTCO1 | CD4+T cell | 0.002 | 0.006 | 0.007 | 0.007 | 0.0055 |
| Macro-c11-CLEC10A | Macropahge | 0.004 | 0.005 | 0.012 | 0.000 | 0.0055 |
| Macro-c15-PCLAF | Macropahge | 0.007 | 0.006 | 0.005 | 0.002 | 0.0049 |
| T-CD4-c10-TNFRSF4 | CD4+T cell | 0.004 | 0.004 | 0.005 | 0.006 | 0.0047 |
| Mono-c01-APOBEC3A | Monocyte | 0.001 | 0.002 | 0.005 | 0.009 | 0.0042 |
| B-c05-CRIP1 | B cell | 0.001 | 0.006 | 0.006 | 0.001 | 0.0036 |
| B-c01-TCL1A | B cell | 0.005 | 0.003 | 0.003 | 0.003 | 0.0035 |
| Epi-c04-CAPS | Epithelial cell | 0.001 | 0.001 | 0.005 | 0.007 | 0.0034 |
| Mono-c02-S100A9 | Monocyte | 0.000 | 0.002 | 0.004 | 0.005 | 0.0028 |
| T-CD4-c03-KLRB1 | CD4+T cell | 0.001 | 0.000 | 0.006 | 0.002 | 0.0025 |
| B-c04-TNFSF9 | B cell | 0.004 | 0.002 | 0.003 | 0.001 | 0.0024 |
| T-CD8-c02-RPL32 | CD8+T cell | 0.004 | 0.003 | 0.001 | 0.001 | 0.0023 |
| Mast-c01-TPSB2 | Mast cell | 0.002 | 0.000 | 0.002 | 0.003 | 0.0018 |
| Neu-c02-IFITM3 | Neutrophil | 0.000 | 0.000 | 0.001 | 0.006 | 0.0018 |
| T-CD4-c04-IFI6 | CD4+T cell | 0.003 | 0.001 | 0.003 | 0.001 | 0.0018 |
| T-CD4-c09-FOXP3 | CD4+T cell | 0.001 | 0.001 | 0.003 | 0.001 | 0.0018 |
| T-CD4-c01-RPL38 | CD4+T cell | 0.002 | 0.003 | 0.002 | 0.001 | 0.0017 |
| Macro-c01-C1QA | Macropahge | 0.000 | 0.006 | 0.001 | 0.000 | 0.0016 |
| cDC-c02-CLEC10A | common dendritic cell | 0.002 | 0.001 | 0.003 | 0.000 | 0.0016 |
| Epi-c03-KIAA1324 | Epithelial cell | 0.001 | 0.000 | 0.001 | 0.005 | 0.0015 |
| T-CD4-c11-PDCD1 | CD4+T cell | 0.002 | 0.001 | 0.002 | 0.001 | 0.0014 |
| NK\|T-c02-FCER1G | Innate lymphoid cells | 0.000 | 0.002 | 0.002 | 0.002 | 0.0014 |
| ILC-c01-SPRY1 | Innate lymphoid cells | 0.001 | 0.001 | 0.001 | 0.002 | 0.0013 |
| Neu-c01-S100A8 | Neutrophil | 0.000 | 0.001 | 0.001 | 0.003 | 0.0013 |
| T-CD8-c08-MALAT1 | CD8+T cell | 0.000 | 0.001 | 0.002 | 0.002 | 0.0013 |
| T-CD4-c02-AREG | CD4+T cell | 0.002 | 0.000 | 0.002 | 0.000 | 0.0012 |
| T-CD8-c01-RPS17 | CD8+T cell | 0.003 | 0.001 | 0.000 | 0.000 | 0.0012 |
| T-CD8-c10-KLRB1 | CD8+T cell | 0.001 | 0.001 | 0.001 | 0.002 | 0.0012 |
| Macro-c14-RPS18 | Macropahge | 0.003 | 0.000 | 0.000 | 0.002 | 0.0011 |
| T-CD8-c07-GZMB | CD8+T cell | 0.001 | 0.001 | 0.001 | 0.001 | 0.0011 |
| T-CD8-c09-LMNA | CD8+T cell | 0.001 | 0.001 | 0.001 | 0.001 | 0.0010 |
| NK\|T-c01-CD3D | Innate lymphoid cells | 0.000 | 0.000 | 0.001 | 0.002 | 0.0010 |
| T-CD4-c06-GZMA | CD4+T cell | 0.000 | 0.000 | 0.002 | 0.001 | 0.0009 |
| T-CD8-c05-CCL4L2 | CD8+T cell | 0.001 | 0.001 | 0.001 | 0.000 | 0.0007 |
| T-CD8-c06-CTLA4 | CD8+T cell | 0.001 | 0.001 | 0.001 | 0.000 | 0.0007 |
| Macro-c13-COL6A3 | Macropahge | 0.001 | 0.000 | 0.002 | 0.000 | 0.0007 |
| Macro-c07-RND3 | Macropahge | 0.001 | 0.001 | 0.001 | 0.000 | 0.0006 |
| T-CD4-c08-TIGIT | CD4+T cell | 0.000 | 0.000 | 0.001 | 0.001 | 0.0006 |
| T-CD8-c04-GZMK | CD8+T cell | 0.001 | 0.000 | 0.000 | 0.001 | 0.0006 |
| Macro-c06-FABP4 | Macropahge | 0.000 | 0.001 | 0.000 | 0.000 | 0.0004 |
| T-CD8-c03-CCL4L2 | CD8+T cell | 0.001 | 0.000 | 0.000 | 0.000 | 0.0004 |
| Macro-c02-ADAMDEC1 | Macropahge | - | - | - | - | - |
| Neu-c03-HIF1AAS3 | Neutrophil | - | - | - | - | - |
| Neu-c04-CXCL10 | Neutrophil | - | - | - | - | - |
| Neu-c05-LTF | Neutrophil | - | - | - | - | - |
| T-CD4-c05-RSAD2 | CD4+T cell | - | - | - | - | - |
| B-c03-IFIT3 | B cell | - | - | - | - | - |
| B-c06-RGS13 | B cell | - | - | - | - | - |
| B-c07-MKI67 | B cell | - | - | - | - | - |

Table S4. Predominant signaling information between cell types is observed in more than half of the pulmonary granulomas.

| **source** | **target** | **pathway_name** | **prob** | **pval** | **rank** |
| --- | --- | --- | --- | --- | --- |
| Macro-c09-CP | Macro-c09-CP | MIF | 0.340474087 | 0 | 1 |
| Macro-c09-CP | Fib-c03-VEGFA | SPP1 | 0.161633075 | 0 | 1 |
| Macro-c09-CP | Macro-c03-APOE | MIF | 0.333020117 | 0 | 1 |
| Macro-c09-CP | Macro-c04-C1S | MIF | 0.353957124 | 0 | 1 |
| Macro-c09-CP | cDC-c01-CLEC9A | MIF | 0.304985738 | 0 | 1 |
| Macro-c09-CP | B-c08-JCHAIN | SPP1 | 0.036602126 | 0 | 1 |
| Macro-c09-CP | T\|B-c01-STMN1 | MIF | 0.278481631 | 0 | 1 |
| Fib-c03-VEGFA | Macro-c09-CP | MIF | 0.390957 | 0 | 1 |
| Fib-c03-VEGFA | Fib-c03-VEGFA | MIF | 0.110723613 | 0 | 1 |
| Fib-c03-VEGFA | Macro-c03-APOE | MIF | 0.382924725 | 0 | 1 |
| Fib-c03-VEGFA | Macro-c04-C1S | MIF | 0.406097739 | 0 | 1 |
| Fib-c03-VEGFA | cDC-c01-CLEC9A | MIF | 0.351349649 | 0 | 1 |
| Fib-c03-VEGFA | B-c08-JCHAIN | ANGPTL | 0.009984161 | 0 | 1 |
| Fib-c03-VEGFA | T\|B-c01-STMN1 | MIF | 0.321778168 | 0 | 1 |
| Macro-c03-APOE | Macro-c09-CP | GALECTIN | 0.13375086 | 0 | 1 |
| Macro-c03-APOE | Fib-c03-VEGFA | GALECTIN | 0.04859665 | 0 | 1 |
| Macro-c03-APOE | Macro-c03-APOE | MIF | 0.117361033 | 0 | 1 |
| Macro-c03-APOE | Macro-c04-C1S | GALECTIN | 0.13231956 | 0 | 1 |
| Macro-c03-APOE | cDC-c01-CLEC9A | MIF | 0.115419626 | 0.03 | 1 |
| Macro-c03-APOE | B-c08-JCHAIN | APRIL | 0.027266506 | 0 | 1 |
| Macro-c03-APOE | T\|B-c01-STMN1 | GALECTIN | 0.106250082 | 0 | 1 |
| Macro-c03-APOE | Fib-c01-CTSK | VISFATIN | 0.049119596 | 0 | 1 |
| Macro-c03-APOE | Fib-c02-CFD | OSM | 0.019045117 | 0 | 1 |
| Macro-c03-APOE | Endo-c01-VWF | CXCL | 0.17972992 | 0 | 1 |
| Macro-c04-C1S | Macro-c09-CP | MIF | 0.136712063 | 0 | 1 |
| Macro-c04-C1S | Fib-c03-VEGFA | VISFATIN | 0.050930062 | 0 | 1 |
| Macro-c04-C1S | Macro-c03-APOE | MIF | 0.123965068 | 0 | 1 |
| Macro-c04-C1S | Macro-c04-C1S | MIF | 0.137435494 | 0 | 1 |
| Macro-c04-C1S | cDC-c01-CLEC9A | MIF | 0.121929508 | 0.01 | 1 |
| Macro-c04-C1S | B-c08-JCHAIN | BAFF | 0.030405604 | 0 | 1 |
| Macro-c04-C1S | T\|B-c01-STMN1 | GALECTIN | 0.10001973 | 0 | 1 |
| Macro-c04-C1S | Fib-c01-CTSK | VISFATIN | 0.054273512 | 0 | 1 |
| Macro-c04-C1S | Fib-c02-CFD | OSM | 0.02530203 | 0 | 1 |
| Macro-c04-C1S | Endo-c01-VWF | VISFATIN | 0.086911015 | 0 | 1 |
| cDC-c01-CLEC9A | Macro-c09-CP | MIF | 0.155663388 | 0 | 1 |
| cDC-c01-CLEC9A | Fib-c03-VEGFA | GALECTIN | 0.043932918 | 0 | 1 |
| cDC-c01-CLEC9A | Macro-c03-APOE | MIF | 0.14143888 | 0 | 1 |
| cDC-c01-CLEC9A | Macro-c04-C1S | MIF | 0.156468926 | 0 | 1 |
| cDC-c01-CLEC9A | cDC-c01-CLEC9A | MIF | 0.139161972 | 0 | 1 |
| cDC-c01-CLEC9A | B-c08-JCHAIN | GALECTIN | 0.016035613 | 0 | 1 |
| cDC-c01-CLEC9A | T\|B-c01-STMN1 | MIF | 0.119289705 | 0 | 1 |
| cDC-c01-CLEC9A | Fib-c01-CTSK | GALECTIN | 0.04376638 | 0 | 1 |
| cDC-c01-CLEC9A | Fib-c02-CFD | GALECTIN | 0.009951262 | 0.02 | 1 |
| cDC-c01-CLEC9A | Endo-c01-VWF | VISFATIN | 0.053976151 | 0 | 1 |
| B-c08-JCHAIN | Macro-c09-CP | MIF | 0.156821002 | 0 | 1 |
| B-c08-JCHAIN | Fib-c03-VEGFA | VISFATIN | 0.003109194 | 0 | 1 |
| B-c08-JCHAIN | Macro-c03-APOE | MIF | 0.255748098 | 0 | 1 |
| B-c08-JCHAIN | Macro-c04-C1S | MIF | 0.272776977 | 0.01 | 1 |
| B-c08-JCHAIN | cDC-c01-CLEC9A | MIF | 0.14021725 | 0 | 1 |
| B-c08-JCHAIN | B-c08-JCHAIN | VISFATIN | 0.000181146 | 0 | 1 |
| B-c08-JCHAIN | T\|B-c01-STMN1 | MIF | 0.120215334 | 0 | 1 |
| B-c08-JCHAIN | Fib-c01-CTSK | VISFATIN | 0.003324302 | 0 | 1 |
| B-c08-JCHAIN | Fib-c02-CFD | GAS | 0.004321021 | 0 | 1 |
| B-c08-JCHAIN | Endo-c01-VWF | VISFATIN | 0.005345733 | 0 | 1 |
| T\|B-c01-STMN1 | Macro-c09-CP | MIF | 0.324099938 | 0 | 1 |
| T\|B-c01-STMN1 | Fib-c03-VEGFA | VISFATIN | 0.021030886 | 0 | 1 |
| T\|B-c01-STMN1 | Macro-c03-APOE | MIF | 0.31686006 | 0 | 1 |
| T\|B-c01-STMN1 | Macro-c04-C1S | MIF | 0.337025006 | 0 | 1 |
| T\|B-c01-STMN1 | cDC-c01-CLEC9A | MIF | 0.290011589 | 0 | 1 |
| T\|B-c01-STMN1 | B-c08-JCHAIN | VISFATIN | 0.001246389 | 0 | 1 |
| T\|B-c01-STMN1 | T\|B-c01-STMN1 | MIF | 0.264551413 | 0 | 1 |
| T\|B-c01-STMN1 | Fib-c01-CTSK | MIF | 0.095940368 | 0.03 | 1 |
| T\|B-c01-STMN1 | Fib-c02-CFD | VISFATIN | 0.006424167 | 0 | 1 |
| T\|B-c01-STMN1 | Endo-c01-VWF | CCL | 0.040536407 | 0 | 1 |
| Fib-c01-CTSK | Macro-c03-APOE | MIF | 0.379590201 | 0 | 1 |
| Fib-c01-CTSK | Macro-c04-C1S | MIF | 0.402620726 | 0 | 1 |
| Fib-c01-CTSK | cDC-c01-CLEC9A | MIF | 0.348245907 | 0 | 1 |
| Fib-c01-CTSK | B-c08-JCHAIN | MK | 0.027528688 | 0 | 1 |
| Fib-c01-CTSK | T\|B-c01-STMN1 | MIF | 0.31887201 | 0 | 1 |
| Fib-c01-CTSK | Fib-c01-CTSK | MIF | 0.116704734 | 0 | 1 |
| Fib-c01-CTSK | Fib-c02-CFD | MK | 0.067562615 | 0 | 1 |
| Fib-c01-CTSK | Endo-c01-VWF | CXCL | 0.085918455 | 0 | 1 |
| Fib-c02-CFD | Macro-c03-APOE | COMPLEMENT | 0.164279673 | 0 | 1 |
| Fib-c02-CFD | Macro-c04-C1S | COMPLEMENT | 0.178195277 | 0 | 1 |
| Fib-c02-CFD | cDC-c01-CLEC9A | COMPLEMENT | 0.060202003 | 0 | 1 |
| Fib-c02-CFD | B-c08-JCHAIN | CXCL | 0.013244251 | 0 | 1 |
| Fib-c02-CFD | T\|B-c01-STMN1 | CXCL | 0.059518086 | 0 | 1 |
| Fib-c02-CFD | Fib-c01-CTSK | VISFATIN | 0.051613282 | 0 | 1 |
| Fib-c02-CFD | Fib-c02-CFD | MK | 0.031777985 | 0 | 1 |
| Fib-c02-CFD | Endo-c01-VWF | VISFATIN | 0.082667444 | 0 | 1 |
| Endo-c01-VWF | Macro-c03-APOE | MIF | 0.105063359 | 0 | 1 |
| Endo-c01-VWF | Macro-c04-C1S | MIF | 0.116753603 | 0 | 1 |
| Endo-c01-VWF | cDC-c01-CLEC9A | MK | 0.021204273 | 0 | 1 |
| Endo-c01-VWF | B-c08-JCHAIN | MK | 0.013907599 | 0 | 1 |
| Endo-c01-VWF | T\|B-c01-STMN1 | MK | 0.031493701 | 0 | 1 |
| Endo-c01-VWF | Fib-c01-CTSK | VISFATIN | 0.040703041 | 0 | 1 |
| Endo-c01-VWF | Fib-c02-CFD | MK | 0.034571663 | 0 | 1 |
| Endo-c01-VWF | Endo-c01-VWF | CCL | 0.066861996 | 0 | 1 |

Table S5. The cell type composition of granulomas in three lymphatic tissue samples.

| Cell subpopulation | Rough classes | Lymphatic granulomas | | | Cell subpopulation average |
| --- | --- | --- | --- | --- | --- |
|  |  | T166353_g1 | T166682_g1 | T166210_g1 |  |
| Macro-c09-CP | Macropahge | 0.540 | 0.538 | 0.168 | 0.415 |
| Fib-c01-CTSK | Fibroblast | 0.031 | 0.068 | 0.010 | 0.037 |
| Macro-c04-C1S | Macropahge | - | - | - | - |
| B-c08-JCHAIN | Plasma cell | 0.011 | 0.009 | 0.050 | 0.023 |
| Macro-c12-IDO1 | Macropahge | - | - | - | - |
| Fib-c03-VEGFA | Fibroblast | 0.011 | 0.016 | 0.017 | 0.015 |
| T\|B-c01-STMN1 | T\|B cells | 0.009 | 0.022 | 0.005 | 0.012 |
| Macro-c03-APOE | Macropahge | 0.001 | 0.009 | 0.056 | 0.022 |
| Endo-c01-VWF | Endothelial cells | 0.005 | 0.014 | 0.072 | 0.030 |
| Fib-c02-CFD | Fibroblast | 0.000 | 0.000 | 0.105 | 0.035 |
| Macro-c10-SPP1 | Macropahge | 0.051 | 0.039 | 0.058 | 0.049 |
| Macro-c05-SELENOP | Macropahge | 0.001 | 0.001 | 0.022 | 0.008 |
| cDC-c01-CLEC9A | common dendritic cell | 0.007 | 0.009 | 0.006 | 0.008 |
| Epi-c01-SFTPC | Epithelial cell | - | - | - | - |
| Macro-c08-EMP1 | Macropahge | - | - | - | - |
| Endo-c02-EDNRB | Endothelial cells | - | - | - | - |
| Mono-c03-CDKN1C | Monocyte | 0.017 | 0.005 | 0.003 | 0.008 |
| B-c02-DUSP4 | B cell | 0.000 | 0.001 | 0.015 | 0.005 |
| T-CD4-c07-CCL4 | CD4+T cell | 0.065 | 0.045 | 0.005 | 0.038 |
| cDC-c03-LAD1 | common dendritic cell | 0.011 | 0.009 | 0.009 | 0.009 |
| Epi-c02-AGER | Epithelial cell | - | - | - | - |
| pDC-c01-GZMB | Plasmacytoid dendritic cells | 0.002 | 0.001 | 0.006 | 0.003 |
| T-CD4-c12-MTCO1 | CD4+T cell | 0.001 | 0.005 | 0.005 | 0.004 |
| Macro-c11-CLEC10A | Macropahge | 0.002 | 0.011 | 0.077 | 0.030 |
| Macro-c15-PCLAF | Macropahge | - | - | - | - |
| T-CD4-c10-TNFRSF4 | CD4+T cell | 0.023 | 0.014 | 0.011 | 0.016 |
| Mono-c01-APOBEC3A | Monocyte | 0.013 | 0.005 | 0.005 | 0.008 |
| B-c05-CRIP1 | B cell | 0.000 | 0.001 | 0.014 | 0.005 |
| B-c01-TCL1A | B cell | 0.000 | 0.000 | 0.053 | 0.018 |
| Epi-c04-CAPS | Epithelial cell | - | - | - | - |
| Mono-c02-S100A9 | Monocyte | 0.003 | 0.004 | 0.002 | 0.003 |
| T-CD4-c03-KLRB1 | CD4+T cell | 0.000 | 0.000 | 0.009 | 0.003 |
| B-c04-TNFSF9 | B cell | 0.000 | 0.000 | 0.012 | 0.004 |
| T-CD8-c02-RPL32 | CD8+T cell | 0.000 | 0.002 | 0.004 | 0.002 |
| Mast-c01-TPSB2 | Mast cell | 0.000 | 0.001 | 0.002 | 0.001 |
| Neu-c02-IFITM3 | Neutrophil | - | - | - | - |
| T-CD4-c04-IFI6 | CD4+T cell | 0.002 | 0.001 | 0.002 | 0.002 |
| T-CD4-c09-FOXP3 | CD4+T cell | 0.001 | 0.001 | 0.008 | 0.003 |
| T-CD4-c01-RPL38 | CD4+T cell | 0.000 | 0.001 | 0.008 | 0.003 |
| Macro-c01-C1QA | Macropahge | 0.075 | 0.110 | 0.070 | 0.085 |
| cDC-c02-CLEC10A | common dendritic cell | 0.006 | 0.003 | 0.009 | 0.006 |
| Epi-c03-KIAA1324 | Epithelial cell | - | - | - | - |
| T-CD4-c11-PDCD1 | CD4+T cell | 0.002 | 0.002 | 0.001 | 0.002 |
| NK\|T-c02-FCER1G | Innate lymphoid cells | 0.003 | 0.001 | 0.002 | 0.002 |
| ILC-c01-SPRY1 | Innate lymphoid cells | 0.001 | 0.001 | 0.001 | 0.001 |
| Neu-c01-S100A8 | Neutrophil | 0.002 | 0.001 | 0.001 | 0.001 |
| T-CD8-c08-MALAT1 | CD8+T cell | 0.000 | 0.000 | 0.016 | 0.005 |
| T-CD4-c02-AREG | CD4+T cell | 0.000 | 0.000 | 0.009 | 0.003 |
| T-CD8-c01-RPS17 | CD8+T cell | 0.000 | 0.001 | 0.008 | 0.003 |
| T-CD8-c10-KLRB1 | CD8+T cell | 0.001 | 0.002 | 0.002 | 0.002 |
| Macro-c14-RPS18 | Macropahge | 0.021 | 0.016 | 0.006 | 0.014 |
| T-CD8-c07-GZMB | CD8+T cell | 0.073 | 0.008 | 0.004 | 0.029 |
| T-CD8-c09-LMNA | CD8+T cell | 0.000 | 0.000 | 0.014 | 0.005 |
| NK\|T-c01-CD3D | Innate lymphoid cells | 0.000 | 0.001 | 0.003 | 0.001 |
| T-CD4-c06-GZMA | CD4+T cell | 0.000 | 0.000 | 0.000 | 0.000 |
| T-CD8-c05-CCL4L2 | CD8+T cell | - | - | - | - |
| T-CD8-c06-CTLA4 | CD8+T cell | 0.001 | 0.003 | 0.001 | 0.001 |
| Macro-c13-COL6A3 | Macropahge | - | - | - | - |
| Macro-c07-RND3 | Macropahge | - | - | - | - |
| T-CD4-c08-TIGIT | CD4+T cell | 0.000 | 0.001 | 0.010 | 0.004 |
| T-CD8-c04-GZMK | CD8+T cell | 0.000 | 0.006 | 0.002 | 0.003 |
| Macro-c06-FABP4 | Macropahge | - | - | - | - |
| T-CD8-c03-CCL4L2 | CD8+T cell | 0.001 | 0.004 | 0.004 | 0.003 |
| Macro-c02-ADAMDEC1 | Macropahge | - | - | - | - |
| Neu-c03-HIF1AAS3 | Neutrophil | - | - | - | - |
| Neu-c04-CXCL10 | Neutrophil | - | - | - | - |
| Neu-c05-LTF | Neutrophil | - | - | - | - |
| T-CD4-c05-RSAD2 | CD4+T cell | 0.002 | 0.002 | 0.000 | 0.001 |
| B-c03-IFIT3 | B cell | 0.000 | 0.000 | 0.008 | 0.003 |
| B-c06-RGS13 | B cell | 0.001 | 0.001 | 0.003 | 0.001 |
| B-c07-MKI67 | B cell | 0.001 | 0.003 | 0.008 | 0.004 |

Table S6. Predominant signaling information between cell types is observed in more than half of the lymphatic granulomas.

| **source** | **target** | **pathway_name** | **prob** | **pval** | **rank** |
| --- | --- | --- | --- | --- | --- |
| Macro-c09-CP | Macro-c09-CP | MIF | 0.227537 | 0 | 1 |
| Macro-c09-CP | Macro-c10-SPP1 | MIF | 0.174463 | 0 | 1 |
| Macro-c09-CP | Macro-c01-C1QA | MIF | 0.125038 | 0 | 1 |
| Macro-c09-CP | Macro-c11-CLEC10A | MIF | 0.377566 | 0 | 1 |
| Macro-c09-CP | T-CD4-c07-CCL4 | MIF | 0.290435 | 0 | 1 |
| Macro-c09-CP | T-CD4-c10-TNFRSF4 | MIF | 0.281904 | 0 | 1 |
| Macro-c09-CP | B-c08-JCHAIN | BAFF | 0.024602 | 0 | 1 |
| Macro-c09-CP | Fib-c01-CTSK | MIF | 0.091888 | 0.03 | 1 |
| Macro-c09-CP | Endo-c01-VWF | CXCL | 0.082965 | 0 | 1 |
| Macro-c10-SPP1 | Macro-c09-CP | MIF | 0.254146 | 0 | 1 |
| Macro-c10-SPP1 | Macro-c10-SPP1 | MIF | 0.196444 | 0 | 1 |
| Macro-c10-SPP1 | Macro-c01-C1QA | MIF | 0.141862 | 0 | 1 |
| Macro-c10-SPP1 | Macro-c11-CLEC10A | MIF | 0.424209 | 0 | 1 |
| Macro-c10-SPP1 | T-CD4-c07-CCL4 | MIF | 0.328486 | 0 | 1 |
| Macro-c10-SPP1 | T-CD4-c10-TNFRSF4 | MIF | 0.319048 | 0 | 1 |
| Macro-c10-SPP1 | B-c08-JCHAIN | MIF | 0.085359 | 0.03 | 1 |
| Macro-c10-SPP1 | Fib-c01-CTSK | MIF | 0.104786 | 0 | 1 |
| Macro-c10-SPP1 | Endo-c01-VWF | CCL | 0.039676 | 0 | 1 |
| Macro-c01-C1QA | Macro-c09-CP | GALECTIN | 0.074619 | 0 | 1 |
| Macro-c01-C1QA | Macro-c10-SPP1 | GALECTIN | 0.024932 | 0 | 1 |
| Macro-c01-C1QA | Macro-c01-C1QA | GALECTIN | 0.009168 | 0.01 | 1 |
| Macro-c01-C1QA | Macro-c11-CLEC10A | GALECTIN | 0.062914 | 0 | 1 |
| Macro-c01-C1QA | T-CD4-c07-CCL4 | CXCL | 0.019174 | 0 | 1 |
| Macro-c01-C1QA | T-CD4-c10-TNFRSF4 | CXCL | 0.016322 | 0 | 1 |
| Macro-c01-C1QA | B-c08-JCHAIN | APRIL | 0.006702 | 0 | 1 |
| Macro-c01-C1QA | Fib-c01-CTSK | GALECTIN | 0.025023 | 0 | 1 |
| Macro-c01-C1QA | Endo-c01-VWF | CCL | 0.009699 | 0 | 1 |
| Macro-c11-CLEC10A | Macro-c09-CP | GALECTIN | 0.174862 | 0 | 1 |
| Macro-c11-CLEC10A | Macro-c10-SPP1 | GALECTIN | 0.058956 | 0 | 1 |
| Macro-c11-CLEC10A | Macro-c01-C1QA | GALECTIN | 0.041197 | 0 | 1 |
| Macro-c11-CLEC10A | Macro-c11-CLEC10A | GALECTIN | 0.148408 | 0 | 1 |
| Macro-c11-CLEC10A | Macro-c14-RPS18 | GALECTIN | 0.016865 | 0.03 | 1 |
| Macro-c11-CLEC10A | T-CD4-c07-CCL4 | CXCL | 0.128899 | 0 | 1 |
| Macro-c11-CLEC10A | T-CD4-c10-TNFRSF4 | CXCL | 0.115753 | 0 | 1 |
| Macro-c11-CLEC10A | B-c08-JCHAIN | BAFF | 0.029363 | 0 | 1 |
| Macro-c11-CLEC10A | Fib-c01-CTSK | GALECTIN | 0.059164 | 0 | 1 |
| Macro-c11-CLEC10A | Endo-c01-VWF | CXCL | 0.135151 | 0 | 1 |
| Macro-c14-RPS18 | Macro-c09-CP | GALECTIN | 0.075344 | 0 | 1 |
| Macro-c14-RPS18 | Macro-c10-SPP1 | GALECTIN | 0.025176 | 0 | 1 |
| Macro-c14-RPS18 | Macro-c01-C1QA | GALECTIN | 0.00926 | 0 | 1 |
| Macro-c14-RPS18 | Macro-c11-CLEC10A | GALECTIN | 0.063528 | 0 | 1 |
| Macro-c14-RPS18 | T-CD4-c07-CCL4 | GALECTIN | 0.066148 | 0 | 1 |
| Macro-c14-RPS18 | T-CD4-c10-TNFRSF4 | GALECTIN | 0.067209 | 0 | 1 |
| Macro-c14-RPS18 | B-c08-JCHAIN | BAFF | 0.007877 | 0 | 1 |
| Macro-c14-RPS18 | Fib-c01-CTSK | GALECTIN | 0.025268 | 0 | 1 |
| Macro-c14-RPS18 | Endo-c01-VWF | MIF | 0.005572 | 0 | 1 |
| T-CD4-c07-CCL4 | Macro-c09-CP | CD40 | 0.084024 | 0 | 1 |
| T-CD4-c07-CCL4 | Macro-c10-SPP1 | CD40 | 0.01976 | 0 | 1 |
| T-CD4-c07-CCL4 | Macro-c01-C1QA | IL16 | 0.005064 | 0 | 1 |
| T-CD4-c07-CCL4 | Macro-c11-CLEC10A | IFN-II | 0.05097 | 0 | 1 |
| T-CD4-c07-CCL4 | T-CD4-c07-CCL4 | MIF | 0.266099 | 0 | 1 |
| T-CD4-c07-CCL4 | T-CD4-c10-TNFRSF4 | MIF | 0.258174 | 0 | 1 |
| T-CD4-c07-CCL4 | B-c08-JCHAIN | LIGHT | 0.002096 | 0 | 1 |
| T-CD4-c07-CCL4 | Fib-c01-CTSK | IFN-II | 0.034842 | 0 | 1 |
| T-CD4-c07-CCL4 | Endo-c01-VWF | CCL | 0.057657 | 0 | 1 |
| T-CD4-c10-TNFRSF4 | Macro-c09-CP | MIF | 0.181992 | 0 | 1 |
| T-CD4-c10-TNFRSF4 | Macro-c10-SPP1 | GALECTIN | 0.009171 | 0 | 1 |
| T-CD4-c10-TNFRSF4 | Macro-c01-C1QA | GALECTIN | 0.005643 | 0 | 1 |
| T-CD4-c10-TNFRSF4 | Macro-c11-CLEC10A | LT | 0.024872 | 0 | 1 |
| T-CD4-c10-TNFRSF4 | Macro-c14-RPS18 | GALECTIN | 0.002302 | 0 | 1 |
| T-CD4-c10-TNFRSF4 | T-CD4-c07-CCL4 | GALECTIN | 0.022112 | 0 | 1 |
| T-CD4-c10-TNFRSF4 | T-CD4-c10-TNFRSF4 | LT | 0.028916 | 0 | 1 |
| T-CD4-c10-TNFRSF4 | B-c08-JCHAIN | CD70 | 0.005636 | 0 | 1 |
| T-CD4-c10-TNFRSF4 | Fib-c01-CTSK | LT | 0.031444 | 0 | 1 |
| T-CD4-c10-TNFRSF4 | Endo-c01-VWF | LT | 0.030571 | 0 | 1 |
| B-c08-JCHAIN | Macro-c09-CP | IL16 | 0.003916 | 0 | 1 |
| B-c08-JCHAIN | Macro-c10-SPP1 | MIF | 0.114534 | 0 | 1 |
| B-c08-JCHAIN | Macro-c01-C1QA | GAS | 0.003049 | 0 | 1 |
| B-c08-JCHAIN | Macro-c11-CLEC10A | IL16 | 0.006163 | 0 | 1 |
| B-c08-JCHAIN | T-CD4-c07-CCL4 | IL16 | 0.003418 | 0 | 1 |
| B-c08-JCHAIN | T-CD4-c10-TNFRSF4 | IL16 | 0.004639 | 0 | 1 |
| B-c08-JCHAIN | Fib-c01-CTSK | GAS | 0.002331 | 0 | 1 |
| B-c08-JCHAIN | Endo-c01-VWF | MIF | 0.006304 | 0 | 1 |
| Fib-c01-CTSK | Macro-c09-CP | COMPLEMENT | 0.239354 | 0 | 1 |
| Fib-c01-CTSK | Macro-c10-SPP1 | COMPLEMENT | 0.041905 | 0 | 1 |
| Fib-c01-CTSK | Macro-c01-C1QA | CSF | 0.012526 | 0 | 1 |
| Fib-c01-CTSK | Macro-c11-CLEC10A | COMPLEMENT | 0.159657 | 0 | 1 |
| Fib-c01-CTSK | Macro-c14-RPS18 | CXCL | 0.038329 | 0 | 1 |
| Fib-c01-CTSK | T-CD4-c07-CCL4 | CXCL | 0.150514 | 0 | 1 |
| Fib-c01-CTSK | T-CD4-c10-TNFRSF4 | CXCL | 0.148724 | 0 | 1 |
| Fib-c01-CTSK | B-c08-JCHAIN | CXCL | 0.042937 | 0 | 1 |
| Fib-c01-CTSK | Fib-c01-CTSK | MIF | 0.0848 | 0 | 1 |
| Fib-c01-CTSK | Endo-c01-VWF | CXCL | 0.07858 | 0 | 1 |
| Endo-c01-VWF | Macro-c09-CP | GALECTIN | 0.185208 | 0 | 1 |
| Endo-c01-VWF | Macro-c10-SPP1 | GALECTIN | 0.062501 | 0 | 1 |
| Endo-c01-VWF | Macro-c01-C1QA | GALECTIN | 0.043782 | 0.01 | 1 |
| Endo-c01-VWF | Macro-c11-CLEC10A | GALECTIN | 0.157295 | 0 | 1 |
| Endo-c01-VWF | Macro-c14-RPS18 | CXCL | 0.004094 | 0 | 1 |
| Endo-c01-VWF | T-CD4-c07-CCL4 | GALECTIN | 0.161779 | 0 | 1 |
| Endo-c01-VWF | T-CD4-c10-TNFRSF4 | GALECTIN | 0.164248 | 0 | 1 |
| Endo-c01-VWF | B-c08-JCHAIN | GALECTIN | 0.013378 | 0 | 1 |
| Endo-c01-VWF | Fib-c01-CTSK | GALECTIN | 0.062721 | 0 | 1 |
| Endo-c01-VWF | Endo-c01-VWF | CCL | 0.093541 | 0 | 1 |

Table S7. The cell type composition of granulomas in three skeletal tissue samples.

| Cell subpopulation | Rough classes | Skeletal granulomas | | | | | Cell subpopulation average |
| --- | --- | --- | --- | --- | --- | --- | --- |
|  |  | T166457_g1 | T166846_g1 | T167126_g1 | T167126_g2 | T167126_g3 |  |
| Macro-c09-CP | Macropahge | 0.040 | 0.056 | 0.010 | 0.029 | 0.008 | 0.028 |
| Fib-c01-CTSK | Fibroblast | 0.089 | 0.068 | 0.079 | 0.042 | 0.048 | 0.065 |
| Macro-c04-C1S | Macropahge | 0.053 | 0.314 | 0.029 | 0.029 | 0.000 | 0.085 |
| B-c08-JCHAIN | Plasma cell | 0.007 | 0.013 | 0.000 | 0.001 | 0.000 | 0.004 |
| Macro-c12-IDO1 | Macropahge | 0.010 | 0.002 | 0.002 | 0.002 | 0.000 | 0.003 |
| Fib-c03-VEGFA | Fibroblast | 0.016 | 0.034 | 0.036 | 0.085 | 0.091 | 0.053 |
| T\|B-c01-STMN1 | T\|B cells | 0.025 | 0.005 | 0.018 | 0.014 | 0.012 | 0.015 |
| Macro-c03-APOE | Macropahge | 0.004 | 0.011 | 0.007 | 0.005 | 0.001 | 0.005 |
| Endo-c01-VWF | Endothelial cells | 0.024 | 0.025 | 0.017 | 0.008 | 0.014 | 0.018 |
| Fib-c02-CFD | Fibroblast | 0.002 | 0.003 | 0.001 | 0.002 | 0.003 | 0.002 |
| Macro-c10-SPP1 | Macropahge | 0.004 | 0.022 | 0.103 | 0.175 | 0.283 | 0.117 |
| Macro-c05-SELENOP | Macropahge | 0.001 | 0.008 | 0.002 | 0.002 | 0.003 | 0.003 |
| cDC-c01-CLEC9A | common dendritic cell | 0.033 | 0.012 | 0.024 | 0.017 | 0.028 | 0.023 |
| Epi-c01-SFTPC | Epithelial cell | - | - | - | - | - | - |
| Macro-c08-EMP1 | Macropahge | 0.000 | 0.000 | 0.000 | 0.006 | 0.001 | 0.002 |
| Endo-c02-EDNRB | Endothelial cells | - | - | - | - | - | - |
| Mono-c03-CDKN1C | Monocyte | 0.000 | 0.000 | 0.000 | 0.001 | 0.000 | 0.000 |
| B-c02-DUSP4 | B cell | 0.006 | 0.000 | 0.001 | 0.001 | 0.002 | 0.002 |
| T-CD4-c07-CCL4 | CD4+T cell | 0.044 | 0.020 | 0.018 | 0.012 | 0.009 | 0.021 |
| cDC-c03-LAD1 | common dendritic cell | 0.036 | 0.029 | 0.012 | 0.006 | 0.010 | 0.019 |
| Epi-c02-AGER | Epithelial cell | - | - | - | - | - | - |
| pDC-c01-GZMB | Plasmacytoid dendritic cells | 0.004 | 0.001 | 0.001 | 0.002 | 0.002 | 0.002 |
| T-CD4-c12-MTCO1 | CD4+T cell | 0.095 | 0.098 | 0.021 | 0.012 | 0.013 | 0.048 |
| Macro-c11-CLEC10A | Macropahge | 0.011 | 0.006 | 0.003 | 0.013 | 0.010 | 0.009 |
| Macro-c15-PCLAF | Macropahge | 0.008 | 0.003 | 0.007 | 0.022 | 0.045 | 0.017 |
| T-CD4-c10-TNFRSF4 | CD4+T cell | 0.015 | 0.010 | 0.012 | 0.011 | 0.008 | 0.011 |
| Mono-c01-APOBEC3A | Monocyte | 0.001 | 0.000 | 0.000 | 0.000 | 0.000 | 0.000 |
| B-c05-CRIP1 | B cell | 0.002 | 0.001 | 0.000 | 0.001 | 0.000 | 0.001 |
| B-c01-TCL1A | B cell | 0.002 | 0.000 | 0.000 | 0.000 | 0.001 | 0.001 |
| Epi-c04-CAPS | Epithelial cell | - | - | - | - | - | - |
| Mono-c02-S100A9 | Monocyte | 0.000 | 0.000 | 0.000 | 0.005 | 0.000 | 0.001 |
| T-CD4-c03-KLRB1 | CD4+T cell | 0.000 | 0.001 | 0.001 | 0.002 | 0.007 | 0.002 |
| B-c04-TNFSF9 | B cell | 0.003 | 0.001 | 0.000 | 0.000 | 0.000 | 0.001 |
| T-CD8-c02-RPL32 | CD8+T cell | 0.001 | 0.001 | 0.001 | 0.002 | 0.001 | 0.001 |
| Mast-c01-TPSB2 | Mast cell | 0.002 | 0.001 | 0.001 | 0.001 | 0.001 | 0.001 |
| Neu-c02-IFITM3 | Neutrophil | 0.000 | 0.000 | 0.000 | 0.000 | 0.000 | 0.000 |
| T-CD4-c04-IFI6 | CD4+T cell | 0.000 | 0.000 | 0.000 | 0.000 | 0.000 | 0.000 |
| T-CD4-c09-FOXP3 | CD4+T cell | 0.002 | 0.001 | 0.001 | 0.002 | 0.000 | 0.001 |
| T-CD4-c01-RPL38 | CD4+T cell | 0.000 | 0.000 | 0.002 | 0.002 | 0.004 | 0.002 |
| Macro-c01-C1QA | Macropahge | 0.001 | 0.004 | 0.087 | 0.044 | 0.016 | 0.031 |
| cDC-c02-CLEC10A | common dendritic cell | 0.011 | 0.004 | 0.009 | 0.007 | 0.004 | 0.007 |
| Epi-c03-KIAA1324 | Epithelial cell | - | - | - | - | - | - |
| T-CD4-c11-PDCD1 | CD4+T cell | 0.001 | 0.005 | 0.002 | 0.002 | 0.004 | 0.003 |
| NK\|T-c02-FCER1G | Innate lymphoid cells | 0.001 | 0.001 | 0.000 | 0.001 | 0.000 | 0.001 |
| ILC-c01-SPRY1 | Innate lymphoid cells | 0.001 | 0.002 | 0.001 | 0.001 | 0.002 | 0.001 |
| Neu-c01-S100A8 | Neutrophil | 0.000 | 0.000 | 0.000 | 0.001 | 0.000 | 0.000 |
| T-CD8-c08-MALAT1 | CD8+T cell | 0.001 | 0.004 | 0.001 | 0.001 | 0.001 | 0.001 |
| T-CD4-c02-AREG | CD4+T cell | 0.000 | 0.001 | 0.000 | 0.001 | 0.004 | 0.001 |
| T-CD8-c01-RPS17 | CD8+T cell | 0.000 | 0.001 | 0.001 | 0.000 | 0.001 | 0.001 |
| T-CD8-c10-KLRB1 | CD8+T cell | 0.000 | 0.002 | 0.000 | 0.000 | 0.001 | 0.001 |
| Macro-c14-RPS18 | Macropahge | 0.000 | 0.000 | 0.000 | 0.001 | 0.001 | 0.000 |
| T-CD8-c07-GZMB | CD8+T cell | 0.001 | 0.002 | 0.000 | 0.001 | 0.000 | 0.001 |
| T-CD8-c09-LMNA | CD8+T cell | 0.001 | 0.002 | 0.001 | 0.001 | 0.002 | 0.001 |
| NK\|T-c01-CD3D | Innate lymphoid cells | 0.000 | 0.000 | 0.000 | 0.000 | 0.000 | 0.000 |
| T-CD4-c06-GZMA | CD4+T cell | 0.003 | 0.007 | 0.003 | 0.004 | 0.004 | 0.004 |
| T-CD8-c05-CCL4L2 | CD8+T cell | - | - | - | - | - | - |
| T-CD8-c06-CTLA4 | CD8+T cell | 0.002 | 0.002 | 0.001 | 0.001 | 0.001 | 0.001 |
| Macro-c13-COL6A3 | Macropahge | 0.001 | 0.000 | 0.000 | 0.000 | 0.000 | 0.000 |
| Macro-c07-RND3 | Macropahge | 0.014 | 0.015 | 0.000 | 0.004 | 0.000 | 0.007 |
| T-CD4-c08-TIGIT | CD4+T cell | 0.002 | 0.004 | 0.001 | 0.002 | 0.002 | 0.002 |
| T-CD8-c04-GZMK | CD8+T cell | 0.000 | 0.000 | 0.001 | 0.000 | 0.001 | 0.001 |
| Macro-c06-FABP4 | Macropahge | 0.050 | 0.004 | 0.051 | 0.020 | 0.002 | 0.025 |
| T-CD8-c03-CCL4L2 | CD8+T cell | 0.001 | 0.002 | 0.001 | 0.000 | 0.000 | 0.001 |
| Macro-c02-ADAMDEC1 | Macropahge | 0.366 | 0.192 | 0.427 | 0.392 | 0.346 | 0.345 |
| Neu-c03-HIF1AAS3 | Neutrophil | 0.002 | 0.002 | 0.001 | 0.001 | 0.002 | 0.002 |
| Neu-c04-CXCL10 | Neutrophil | 0.000 | 0.000 | 0.000 | 0.000 | 0.000 | 0.000 |
| Neu-c05-LTF | Neutrophil | 0.000 | 0.000 | 0.000 | 0.002 | 0.001 | 0.001 |
| T-CD4-c05-RSAD2 | CD4+T cell | 0.000 | 0.000 | 0.000 | 0.003 | 0.002 | 0.001 |
| B-c03-IFIT3 | B cell | 0.000 | 0.000 | 0.000 | 0.000 | 0.000 | 0.000 |
| B-c06-RGS13 | B cell | - | - | - | - | - | - |
| B-c07-MKI67 | B cell | - | - | - | - | - | - |

Table S8. Predominant signaling information between cell types is observed in more than half of the skeletal granulomas.

| **source** | **target** | **pathway_name** | **prob** | **pval** | **rank** |
| --- | --- | --- | --- | --- | --- |
| Macro-c09-CP | Macro-c09-CP | MIF | 0.188404 | 0 | 1 |
| Macro-c09-CP | Macro-c10-SPP1 | MIF | 0.177726 | 0 | 1 |
| Macro-c09-CP | Fib-c03-VEGFA | SPP1 | 0.203441 | 0 | 1 |
| Macro-c09-CP | Macro-c02-ADAMDEC1 | MIF | 0.334726 | 0 | 1 |
| Macro-c09-CP | cDC-c01-CLEC9A | MIF | 0.154343 | 0 | 1 |
| Macro-c09-CP | cDC-c03-LAD1 | MIF | 0.302794 | 0 | 1 |
| Macro-c09-CP | T-CD4-c07-CCL4 | MIF | 0.275423 | 0 | 1 |
| Macro-c09-CP | T-CD4-c12-MTCO1 | MIF | 0.257774 | 0 | 1 |
| Macro-c09-CP | T\|B-c01-STMN1 | MIF | 0.272617 | 0 | 1 |
| Macro-c10-SPP1 | Macro-c09-CP | MIF | 0.378696 | 0 | 1 |
| Macro-c10-SPP1 | Macro-c10-SPP1 | MIF | 0.2101 | 0 | 1 |
| Macro-c10-SPP1 | Fib-c03-VEGFA | SPP1 | 0.420899 | 0 | 1 |
| Macro-c10-SPP1 | Macro-c02-ADAMDEC1 | MIF | 0.396548 | 0 | 1 |
| Macro-c10-SPP1 | cDC-c01-CLEC9A | MIF | 0.316677 | 0 | 1 |
| Macro-c10-SPP1 | cDC-c03-LAD1 | MIF | 0.359869 | 0 | 1 |
| Macro-c10-SPP1 | T-CD4-c07-CCL4 | MIF | 0.328409 | 0 | 1 |
| Macro-c10-SPP1 | T-CD4-c12-MTCO1 | MIF | 0.308043 | 0 | 1 |
| Macro-c10-SPP1 | T\|B-c01-STMN1 | MIF | 0.325124 | 0 | 1 |
| Fib-c03-VEGFA | Macro-c09-CP | MIF | 0.325673 | 0.015 | 1 |
| Fib-c03-VEGFA | Macro-c10-SPP1 | MIF | 0.181152 | 0 | 1 |
| Fib-c03-VEGFA | Fib-c03-VEGFA | MIF | 0.11065 | 0.03 | 1 |
| Fib-c03-VEGFA | Macro-c02-ADAMDEC1 | MIF | 0.341256 | 0 | 1 |
| Fib-c03-VEGFA | cDC-c01-CLEC9A | MIF | 0.270792 | 0.015 | 1 |
| Fib-c03-VEGFA | cDC-c03-LAD1 | MIF | 0.308805 | 0 | 1 |
| Fib-c03-VEGFA | T-CD4-c07-CCL4 | MIF | 0.280988 | 0 | 1 |
| Fib-c03-VEGFA | T-CD4-c12-MTCO1 | MIF | 0.263043 | 0 | 1 |
| Fib-c03-VEGFA | T\|B-c01-STMN1 | MIF | 0.278131 | 0 | 1 |
| Macro-c02-ADAMDEC1 | Macro-c09-CP | GALECTIN | 0.1358 | 0 | 1 |
| Macro-c02-ADAMDEC1 | Macro-c10-SPP1 | GALECTIN | 0.090601 | 0 | 1 |
| Macro-c02-ADAMDEC1 | Fib-c03-VEGFA | GALECTIN | 0.067629 | 0 | 1 |
| Macro-c02-ADAMDEC1 | Macro-c02-ADAMDEC1 | GALECTIN | 0.133744 | 0 | 1 |
| Macro-c02-ADAMDEC1 | cDC-c01-CLEC9A | GALECTIN | 0.08814 | 0 | 1 |
| Macro-c02-ADAMDEC1 | cDC-c03-LAD1 | GALECTIN | 0.06612 | 0 | 1 |
| Macro-c02-ADAMDEC1 | T-CD4-c07-CCL4 | GALECTIN | 0.120361 | 0 | 1 |
| Macro-c02-ADAMDEC1 | T-CD4-c12-MTCO1 | GALECTIN | 0.127096 | 0 | 1 |
| Macro-c02-ADAMDEC1 | T\|B-c01-STMN1 | GALECTIN | 0.108556 | 0 | 1 |
| Macro-c02-ADAMDEC1 | Fib-c01-CTSK | GALECTIN | 0.049652 | 0 | 1 |
| Macro-c02-ADAMDEC1 | Endo-c01-VWF | CXCL | 0.267353 | 0.001667 | 1 |
| cDC-c01-CLEC9A | Macro-c09-CP | MIF | 0.160074 | 0 | 1 |
| cDC-c01-CLEC9A | Macro-c10-SPP1 | MIF | 0.150704 | 0 | 1 |
| cDC-c01-CLEC9A | Fib-c03-VEGFA | GALECTIN | 0.057381 | 0 | 1 |
| cDC-c01-CLEC9A | Macro-c02-ADAMDEC1 | MIF | 0.151294 | 0 | 1 |
| cDC-c01-CLEC9A | cDC-c01-CLEC9A | GALECTIN | 0.074428 | 0 | 1 |
| cDC-c01-CLEC9A | cDC-c03-LAD1 | MIF | 0.144173 | 0 | 1 |
| cDC-c01-CLEC9A | T-CD4-c07-CCL4 | GALECTIN | 0.102088 | 0 | 1 |
| cDC-c01-CLEC9A | T-CD4-c12-MTCO1 | GALECTIN | 0.107888 | 0 | 1 |
| cDC-c01-CLEC9A | T\|B-c01-STMN1 | GALECTIN | 0.091991 | 0 | 1 |
| cDC-c01-CLEC9A | Fib-c01-CTSK | VISFATIN | 0.0477 | 0 | 1 |
| cDC-c01-CLEC9A | Endo-c01-VWF | CXCL | 0.169888 | 0 | 1 |
| cDC-c03-LAD1 | Macro-c09-CP | MIF | 0.128968 | 0 | 1 |
| cDC-c03-LAD1 | Macro-c10-SPP1 | MIF | 0.121155 | 0 | 1 |
| cDC-c03-LAD1 | Fib-c03-VEGFA | VISFATIN | 0.058735 | 0 | 1 |
| cDC-c03-LAD1 | Macro-c02-ADAMDEC1 | MIF | 0.121646 | 0 | 1 |
| cDC-c03-LAD1 | cDC-c01-CLEC9A | GALECTIN | 0.074279 | 0 | 1 |
| cDC-c03-LAD1 | cDC-c03-LAD1 | CCL | 0.200775 | 0 | 1 |
| cDC-c03-LAD1 | T-CD4-c07-CCL4 | CXCL | 0.13054 | 0 | 1 |
| cDC-c03-LAD1 | T-CD4-c12-MTCO1 | GALECTIN | 0.107678 | 0 | 1 |
| cDC-c03-LAD1 | T\|B-c01-STMN1 | GALECTIN | 0.09181 | 0 | 1 |
| cDC-c03-LAD1 | Fib-c01-CTSK | VISFATIN | 0.05063 | 0 | 1 |
| cDC-c03-LAD1 | Endo-c01-VWF | CXCL | 0.382895 | 0 | 1 |
| T-CD4-c07-CCL4 | Macro-c09-CP | MIF | 0.17928 | 0 | 1 |
| T-CD4-c07-CCL4 | Macro-c10-SPP1 | MIF | 0.169012 | 0 | 1 |
| T-CD4-c07-CCL4 | Fib-c03-VEGFA | VISFATIN | 0.034804 | 0 | 1 |
| T-CD4-c07-CCL4 | Macro-c02-ADAMDEC1 | MIF | 0.318131 | 0 | 1 |
| T-CD4-c07-CCL4 | cDC-c01-CLEC9A | MIF | 0.146571 | 0 | 1 |
| T-CD4-c07-CCL4 | cDC-c03-LAD1 | MIF | 0.287537 | 0 | 1 |
| T-CD4-c07-CCL4 | T-CD4-c07-CCL4 | MIF | 0.261316 | 0 | 1 |
| T-CD4-c07-CCL4 | T-CD4-c12-MTCO1 | MIF | 0.116242 | 0 | 1 |
| T-CD4-c07-CCL4 | T\|B-c01-STMN1 | MIF | 0.258642 | 0 | 1 |
| T-CD4-c07-CCL4 | Fib-c01-CTSK | IFN-II | 0.037331 | 0 | 1 |
| T-CD4-c07-CCL4 | Endo-c01-VWF | CCL | 0.135229 | 0 | 1 |
| T-CD4-c12-MTCO1 | Macro-c09-CP | MIF | 0.119988 | 0 | 1 |
| T-CD4-c12-MTCO1 | Macro-c10-SPP1 | MIF | 0.11265 | 0.03 | 1 |
| T-CD4-c12-MTCO1 | Fib-c03-VEGFA | VISFATIN | 0.018341 | 0 | 1 |
| T-CD4-c12-MTCO1 | Macro-c02-ADAMDEC1 | MIF | 0.113111 | 0.01 | 1 |
| T-CD4-c12-MTCO1 | cDC-c01-CLEC9A | IFN-II | 0.012453 | 0 | 1 |
| T-CD4-c12-MTCO1 | cDC-c03-LAD1 | IFN-II | 0.023962 | 0 | 1 |
| T-CD4-c12-MTCO1 | T-CD4-c07-CCL4 | CCL | 0.023407 | 0 | 1 |
| T-CD4-c12-MTCO1 | T-CD4-c12-MTCO1 | TNF | 0.007913 | 0 | 1 |
| T-CD4-c12-MTCO1 | T\|B-c01-STMN1 | TNF | 0.01653 | 0 | 1 |
| T-CD4-c12-MTCO1 | Fib-c01-CTSK | IFN-II | 0.024094 | 0 | 1 |
| T-CD4-c12-MTCO1 | Endo-c01-VWF | CCL | 0.087821 | 0 | 1 |
| T\|B-c01-STMN1 | Macro-c09-CP | MIF | 0.179098 | 0 | 1 |
| T\|B-c01-STMN1 | Macro-c10-SPP1 | MIF | 0.168837 | 0 | 1 |
| T\|B-c01-STMN1 | Fib-c03-VEGFA | VISFATIN | 0.036497 | 0 | 1 |
| T\|B-c01-STMN1 | Macro-c02-ADAMDEC1 | MIF | 0.317799 | 0 | 1 |
| T\|B-c01-STMN1 | cDC-c01-CLEC9A | MIF | 0.146416 | 0.01 | 1 |
| T\|B-c01-STMN1 | cDC-c03-LAD1 | MIF | 0.287232 | 0 | 1 |
| T\|B-c01-STMN1 | T-CD4-c07-CCL4 | MIF | 0.261035 | 0 | 1 |
| T\|B-c01-STMN1 | T-CD4-c12-MTCO1 | MIF | 0.116114 | 0 | 1 |
| T\|B-c01-STMN1 | T\|B-c01-STMN1 | MIF | 0.258363 | 0 | 1 |
| T\|B-c01-STMN1 | Fib-c01-CTSK | VISFATIN | 0.031358 | 0 | 1 |
| T\|B-c01-STMN1 | Endo-c01-VWF | CCL | 0.101216 | 0 | 1 |
| Fib-c01-CTSK | Macro-c02-ADAMDEC1 | MIF | 0.345092 | 0 | 1 |
| Fib-c01-CTSK | cDC-c01-CLEC9A | MIF | 0.273956 | 0 | 1 |
| Fib-c01-CTSK | cDC-c03-LAD1 | MIF | 0.312338 | 0 | 1 |
| Fib-c01-CTSK | T-CD4-c07-CCL4 | MIF | 0.28426 | 0 | 1 |
| Fib-c01-CTSK | T-CD4-c12-MTCO1 | MIF | 0.266142 | 0 | 1 |
| Fib-c01-CTSK | T\|B-c01-STMN1 | MIF | 0.281373 | 0 | 1 |
| Fib-c01-CTSK | Fib-c01-CTSK | MK | 0.067998 | 0 | 1 |
| Fib-c01-CTSK | Endo-c01-VWF | CXCL | 0.394218 | 0 | 1 |
| Endo-c01-VWF | Macro-c02-ADAMDEC1 | MIF | 0.147344 | 0 | 1 |
| Endo-c01-VWF | cDC-c01-CLEC9A | MK | 0.018668 | 0 | 1 |
| Endo-c01-VWF | cDC-c03-LAD1 | MIF | 0.140378 | 0.02 | 1 |
| Endo-c01-VWF | T-CD4-c07-CCL4 | CXCL | 0.042862 | 0 | 1 |
| Endo-c01-VWF | T-CD4-c12-MTCO1 | CXCL | 0.021893 | 0 | 1 |
| Endo-c01-VWF | T\|B-c01-STMN1 | CXCL | 0.036705 | 0 | 1 |
| Endo-c01-VWF | Fib-c01-CTSK | MK | 0.03308 | 0 | 1 |
| Endo-c01-VWF | Endo-c01-VWF | CCL | 0.178487 | 0 | 1 |

### Supplemental Data files

Data file S1**.** **Top 20 up-regulated and down-regulated expressed genes in each granuloma compared to non-granulomatous regions in the same sample.**
